# The Receptor Kinase FER Mediates Phase Separation of Glycine-Rich RNA-Binding Protein 7 to Confer Temperature Resilience in *Arabidopsis*

**DOI:** 10.1101/2022.03.06.483201

**Authors:** Fan Xu, Long Wang, Yingbin Li, Junfeng Shi, Dorothee Staiger, Weijun Chen, Lifeng Wang, Feng Yu

## Abstract

Temperature fluctuations repress plant growth. Although glycine-rich RNA-binding proteins (GRPs) and cold shock proteins (CSPs) have been implicated in cold adaptation, their physiological roles in the response to temperature fluctuations are largely unknown. The receptor kinase FERONIA (FER), a master regulator of cell growth, phosphorylates GRP7 within its intrinsically disordered region to modulate mRNA alternative splicing in the nucleus. Here we show that natural variations at a GRP7 residue phosphorylated by FER influences GRP7 liquid-liquid phase separation (LLPS), aiding *Arabidopsis* to grow over a wider temperature range. LLPS of GRP7 in the cytoplasm leads to the formation of stress granules that recruits RNAs, along with the translation machinery component eIF4E1 and mRNA chaperones, CSP1 and CSP3, to inhibit translation. Mutations in FER and the GRP7-LLPS-recruited components attenuate root growth under temperature shift conditions. Our findings illustrate the roles of GRP7 LLPS in improving plant root capacity to withstand temperature fluctuations.

## Introduction

Temperature fluctuations profoundly influence plant growth and development^1, 2^. In many regions, periodic differences between minimum and maximum temperatures result in high temperature fluctuations. Plants have evolved multiple protective mechanisms to cope with these chronic changes^3^.

Low environmental temperatures (cold stress) are associated with phenotypic symptoms in aerial tissues including leaf yellowing (chlorosis) and reduced leaf expansion and wilting^4^. Plants cope with cold stress by altering their gene expression program for protection against damage caused by low temperatures^1-3^. Over the past three decades, many cold-regulated genes and their regulatory components have been identified (reviewed in ref. ^2, 3, 5^). Members of the glycine-rich RNA-binding protein (GRP) and cold shock domain protein (CSP) families accumulate in response to cold stress and play essential roles in the post-transcriptional regulation of gene expression^6-12^. For example, GRP7 contains an N-terminal RNA recognition motif and a C-terminal glycine-rich domain: the N-terminal RNA recognition motif contributes to its nucleic-acid-binding and RNA chaperone activity, while the C-terminal glycine-rich region contributes to its attaining full activity by an unknown mechanism^13^. Heterologous expression experiments with plant GRP7 in the highly cold-sensitive *Escherichia coli* mutant BX04 showed that GRP7 acts as an RNA chaperone that destabilizes the secondary structures of RNA molecules to facilitate their digestion into smaller fragments^8^. *In planta*, overexpression of Arabidopsis (*Arabidopsis thaliana*) *CSP3* or *GRP7* in transgenic plants conferred enhanced freezing tolerance relative to wild-type plants^9, 10^. GRP7 was thus proposed to directly or indirectly transport RNA from the nucleus to the cytoplasm^9^. However, whether and how GRP7, CSP1 and/or CSP3 impart an elevated tolerance of temperature fluctuation to plants is far from resolved.

The glycine-rich domain of GRP7 is a typical intrinsically disordered region (IDR). IDRs effectively trigger liquid-liquid phase separation (LLPS)^14-16^. In plants, LLPS is reported to be involved in sensing high temperature^17^, water potential^18^, phytohormone^19^, and pathogens^20, 21^. At the molecular level, LLPS is driven by weak, transient interactions between proteins or other molecules and multivalent domains or IDRs^14, 15^. LLPS can be regulated by various post-translational modifications, such as phosphorylation, of its driving proteins^22^. In an *in vitro* reconstituted system, phosphorylation affected the assembly of peptide-RNA liquid droplets^23^. In living cells, an alteration of interaction strength due to protein phosphorylation can manifest in several distinct ways, including enhanced internal dynamics^24^, protein release from condensates^25^, and dissolution of the entire condensate structure^26^. Phosphorylation can also stimulate LLPS when specific protein-protein interactions are involved, such as the binding of phosphorylated Src homology 2 on a specific tyrosine residue and its interacting partners in kidney podocytes^27^. In addition, LLPS might underlie the formation of many different condensates^14^, which is relevant to stress granules (SGs) formation^28^. SGs are transient cytoplasmic condensates composed of mRNAs that are stalled in translation initiation and mRNA ribonucleoproteins (mRNPs). SGs are highly conserved, being found in plants, mammals, and yeast (*Saccharomyces cerevisiae*) (reviewed in ref.^29^). Sequestration of mRNAs into SGs provides a dynamic mechanism to repress translational activity during a stress response, while allowing the expeditious resumption of translational activity after the stress passes^30^. However, how temperature fluctuations might translate into LLPS modulation in plants has been largely unknown.

Complex signalling cascades are initiated by receptors on the plasma membrane in response to environmental or endogenous cues. Receptor-like kinases (RLKs; or receptor kinases when their ligands are known) initiate signalling cascades that target various stress response modules. Over 600 *RLK* genes have been identified in Arabidopsis^31, 32^. One such RLKs, FERONIA (FER, from the *Catharanthus roseus* receptor-like kinase 1-like [*Cr*RLKL1] RLK subfamily), participates in cold responses^33-35^. For example, perceiving the low temperature by FER controls Arabidopsis root hair growth^33, 36^. We previously showed that upon activation by its ligand RAPID ALKALINIZATION FACTOR 1 (RALF1), FER phosphorylates GRP7, altering its RNA-binding properties and association with the spliceosome component U1-70K, which triggers rapid, global changes in alternative splicing^37^.

Here, we discovered that natural variations at a FER phosphorylation site within GRP7 affects its LLPS, thereby facilitating root growth under harsh temperature changes. Cytoplasmic GRP7 LLPS leads to the formation of SGs, which facilitates RNA assembly and sequestration of the translation initiation factor eIF4E1 and the mRNA chaperones CSP1 and CSP3, and subsequently blocking translation. Our study provides a novel example of plant cell signalling pathway-specific LLPS that confers an adaptation to temperature fluctuation.

## Results

### A FER-dependent phosphorylation site within the intrinsically disordered region of GRP7 is required for adaptation to high temperature fluctuation and is under natural selection

*GRP7* expression is regulated by cold^8, 9^, but how GRP7 then directs temperature response is still largely unknown. We first confirmed that GRP7 promotes cold (16 °C) adaptation in the *E. coli* strain BL21. Bacteria heterologously producing a GRP7-GFP (green fluorescent protein) fusion protein grew faster than those producing GFP alone starting 36h after transfer to 16 °C (Fig. 1a, ref.^8^). Given that phosphorylation of RNA-binding proteins profoundly impacts their activities/functions, and that GRP7 carries six FER-dependent phosphorylated residues (Tyr111, Ser112, Ser132, Tyr138, Ser139, and Ser140) (Fig. 1b, ref.^37^), we then searched for single-nucleotide polymorphisms (SNPs) in GRP7 among 1,135 Arabidopsis accessions^38^. We identified SNPs in the accessions Blh-1 and BRE-14, which lead to the substitution of Ser132 to Ala (Blh-1) or Tyr131 to Cys and Ser132 to Gly (BRE-14). Notably, both original residues in Blh-1 and BRE-14 were potential phosphorylation sites within the IDR (Fig. 1b). Compared to the wild-type accession Col-0, Blh-1 and BRE-14 experience a smaller range of temperatures (*T*_*max*_ - *T*_*min*_, Fig.1c) at their original collection sites, suggesting that this FER-dependent phosphorylation site (Ser132) of GRP7 phase separation may be under natural selection. To test whether this site contributes to seedling adaptation to temperature fluctuations, we generated transgenic lines in the *grp7-1 8i* background (a *grp7-1* mutant with GRP8 silence by RNA interference) that expressed in the GRP7 variant from Blh-1 and BRE-14 under the control of GRP7 promoter. We grew these seedlings under daily fluctuating temperature conditions, ranging from 10 °C to 34 °C in intervals of 2 °C, for 5 d (*T*_*max*_ - *T*_*min*_ = 24°C, see Fig. 1d, top). We found that the *grp7-1 8i* genotype was sensitive to fluctuating temperature and grew poorly, as indicated by shorter primary roots. Whereas plants with the Col-0 version of GRP7 showed normal primary root growth under the same conditions. Importantly, neither the Blh-1 version (lines *pGRP7::GRP7*^*mut132S(A)*^*-GFP* #11 and #12) nor the BRE-14 version (lines *pGRP7::GRP7*^*mut131Y(C)132S(G)*^*-GFP* #14 and #15) restored the temperature sensitivity of *grp7-1 8i* relative to *pGRP7::GRP7-GFP* (Fig. 1d). Collectively, these results suggested that the phosphorylation of Ser132 in GRP7 is under natural selection and modulates the response to temperature fluctuations in Arabidopsis.

**Fig. 1.**
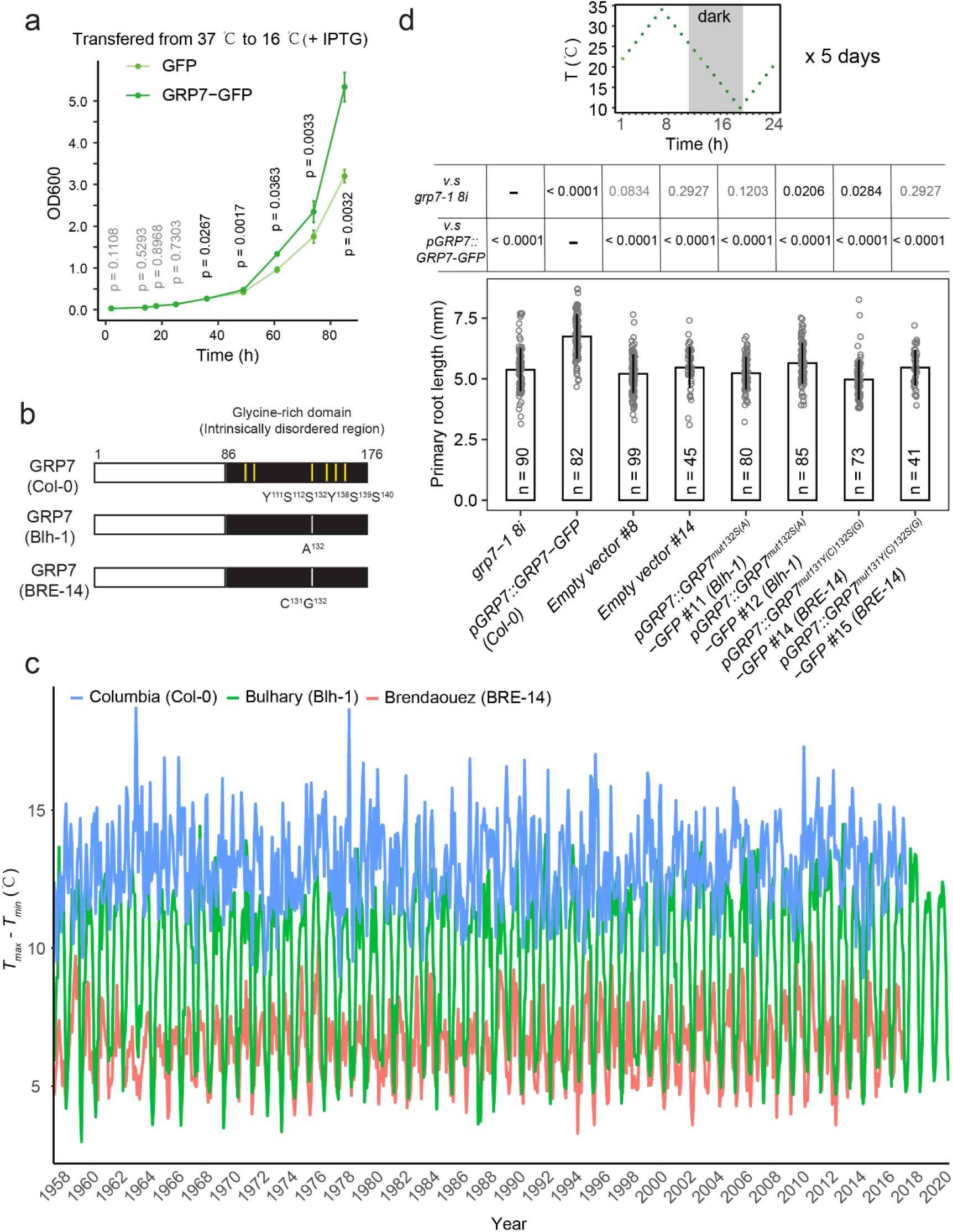
A FER-dependent phosphorylation site in GRP7 contributes to Arabidopsis seedling adaptation to a wider temperature range. **a**, Growth curve of *Escherichia. coli* BL21 strains expressing *GRP7-GFP* (green fluorescent protein) or GFP after transferred from 37 °C to 16 °C over 84 h. The production of recombinant proteins was induced by the addition of isopropyl-β-D-thiogalactopyranoside (IPTG). **b**, Schematic representation of natural variations in GRP7. Vertical yellow bars indicate FER phosphorylation sites indicated according to Wang et al (2020); grey bars indicate variants in the Blh-1 and BRE-14 accessions. **c**, Monthly average temperature range (= highest [T_max_] - lowest [T_min_]) at the original collection sites of the indicated accessions. **d**, Primary root length of *GRP7*-related materials grown under fluctuating temperature with *T*_*max*_ - *T*_*min*_ = 24 °C. Three -day-old seedlings were grown under the temperature regime illustrated in the top panel for 5 d. Empty vector or genomic constructs expressing the GRP7 variants from accessions Blh-1 and BRE-14 under the control of the GRP7 promoter were transformed into *grp7-1 8i*. Two independent lines were selected for each construct. For **a**,**d**, the data are presented as the means ±standard deviation (s.d.) values. *P* values were calculated by one-way ANOVA (middle table).

### The intrinsically disordered region of GRP7 drives its liquid-liquid phase separation

To dissect the molecular mechanism by which Ser132 contributes to GRP7 function, we characterized the C-terminal IDR within GRP7. The IDR of GRP7 was enriched in low-complexity polyglycine (Supplementary Fig. 1a) and showed a high score for a prion-like domain (PrLD; Fig. 2a). PrLDs allow the formation of weak protein-protein interactions that facilitate the development of a dynamic, multivalent meshworks manifesting as phase-separated droplets^14, 15^. This finding prompted us to investigate whether GRP7 condensates have liquid-like characteristics in living root cells of transgenic seedlings harbouring the *pGRP7::GRP7-GFP* construct. Indeed, we observed bright cytoplasmic condensates upon treating seedlings with the RALF1 peptide, a ligand of FER (Fig. 2b). To investigate the formation of GRP7-GFP condensates upon activation of FER, we treated 5-d-old *GRP7-GFP* seedlings with 2 μM RALF1 peptide for various times (0-8 h). After 5 h of treatment, we detected spherical GRP7-GFP condensates within both the cytoplasm and nucleoplasm of whole root tissues, especially for the cells in the transition zone (TZ; Fig. 2b,c, Movie 1). Protoplasts isolated from young *GRP7-GFP* roots were isolated and treated with RALF1 for 2h showed the same pattern of GRP7-GFP condensate (Fig. 2d). We detected no condensates were detected in root cells or their protoplasts expressing GFP alone (Fig. 2b-d). The cytoplasmic GRP7-GFP condensates varied in size from ∼0.5 μm to 2.0 μm (> 95% of total) (Fig. 2e), within the size range of typical intracellular condensates^15^. We also conducted fluorescence recovery after photobleaching (FRAP) was to circumvent artifacts caused by random motion. We observed the rapid redistribution of GRP7-GFP from the unbleached area to the bleached area (Fig. 2f), indicating that GRP7-GFP molecules diffuse in and out of condensates. GRP7-GFP condensates fused and relaxed into a single body as soon as they intersected (Fig. 2g, Movie 2), which is a typical characteristic of LLPS^15^.

**Fig. 2.**
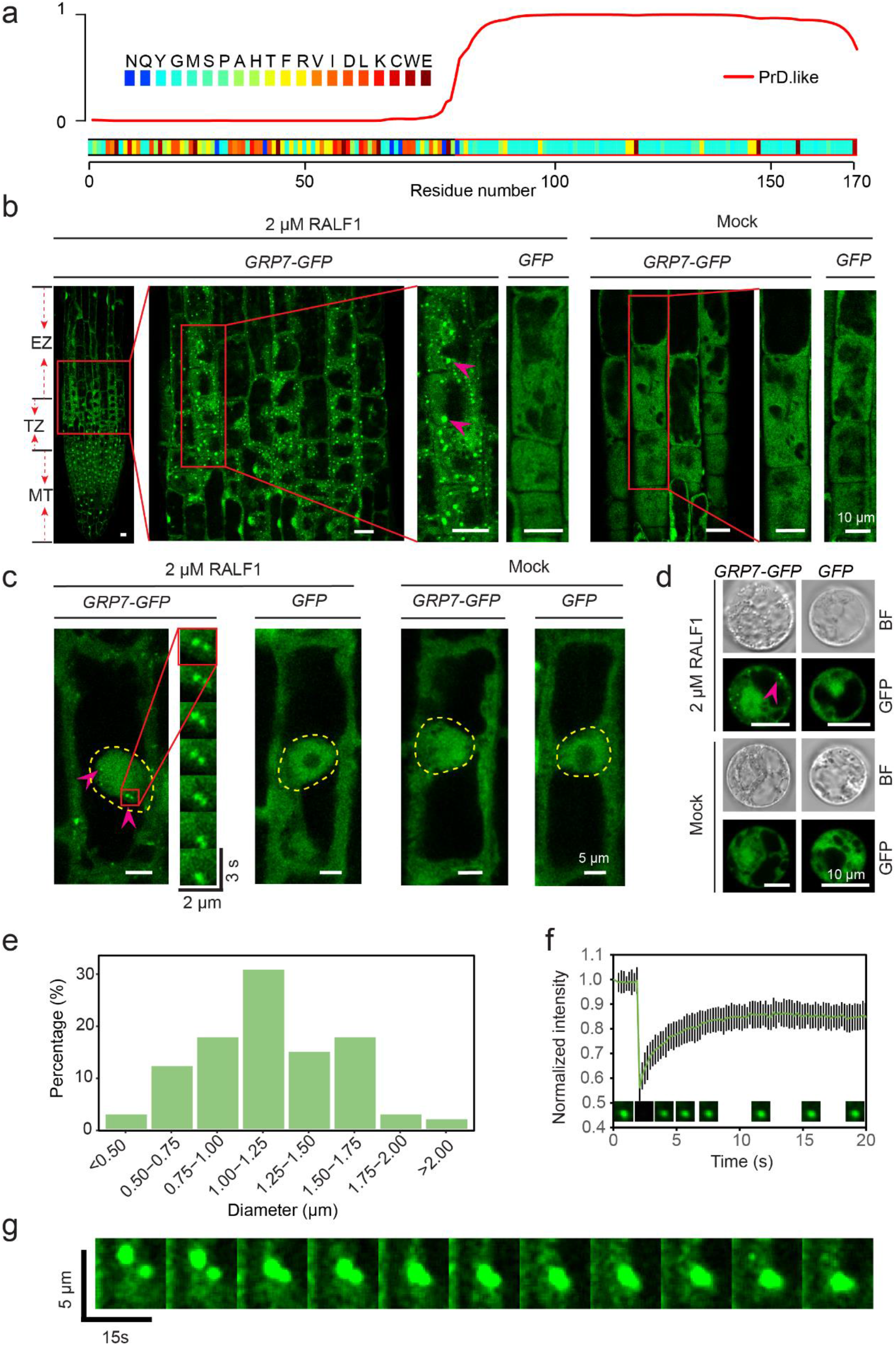
GRP7 undergoes phase separation in *Arabidopsis* root cells. Predictions of prion-like domains (PrLDs) in GRP7 with the online tool Prion-like Amino Acid Composition (PLAAC; http://plaac.wi.mit.edu/). Abbreviations for amino acids: A, Ala; C, Cys; D, Asp; E, Glu; F, Phe; G, Gly; H, His; I, Ile; K, Lys; L, Leu; M, Met; N, Asn; P, Pro; Q, Gln; R, Arg; S, Ser; T, Thr; V, Val; W, Trp; Y, Tyr. **b**,**c**, GRP7 forms condensates in the cytoplasm (**b**) and the nucleoplasm (**c**) in root cells of 5-d-old seedlings treated with RALF1 for 5h. EZ, elongation zone, TZ, transition zone, MT, meristem. Representative data from five independent experiments. Magenta arrowheads point toward GRP7 condensates. The time-lapse in **c** shows the fusion of two nucleoplasmic GRP7 foci. The dashed yellow lines enclose the nucleus. **d**, GRP7 condensates in root cell protoplasts treated with RALF1 for 5h. Magenta arrowhead points to GRP7 condensates. **e**, Size distribution of GRP7 condensates. n = 108 droplets from five seedlings. **f**, Fluorescence intensity during FRAP of GRP7 droplets. Data are shown as the means ± s.d. values (n = 9). The intensities were normalized to the pre-bleached levels, set to 1.0. **g**, Fusion of GRP7 cytoplasmic foci in root cells of 5-d-old seedlings treated with RALF1 for 5 h. Representative data from five independent experiments.

GRP7-GFP condensates formed quickly upon heat exposure. Indeed, we detected GRP7 condensates in root cells as early as 15-30 min after a shift from 22 °C to 38 °C, but not after a shift from 22 °C to 4 °C (Supplementary Fig. 2a,b). In addition, the heat-induced formation of GRP7-GFP condensates was reversible (Supplementary Fig. 2b), suggesting that their formation could be an adaptation to the temperature changes.

Several membraneless condensates have been reported to exist in plant cells, including the nucleolus, Cajal bodies, nuclear speckles, DNA damage foci, dicing bodies, and photobodies in the nucleus and processing bodies (PBs) and stress granules (SGs) in the cytoplasm (reviewed in ref.^30^). With various pharmacological treatments, we established that the cytoplasmic GRP7 condensates are SGs (see Supplementary Text, Supplementary Figs. 2 and 3). Collectively, these results provided experimental evidence that GRP7 undergoes LLPS in root cells.

**Fig. 3.**
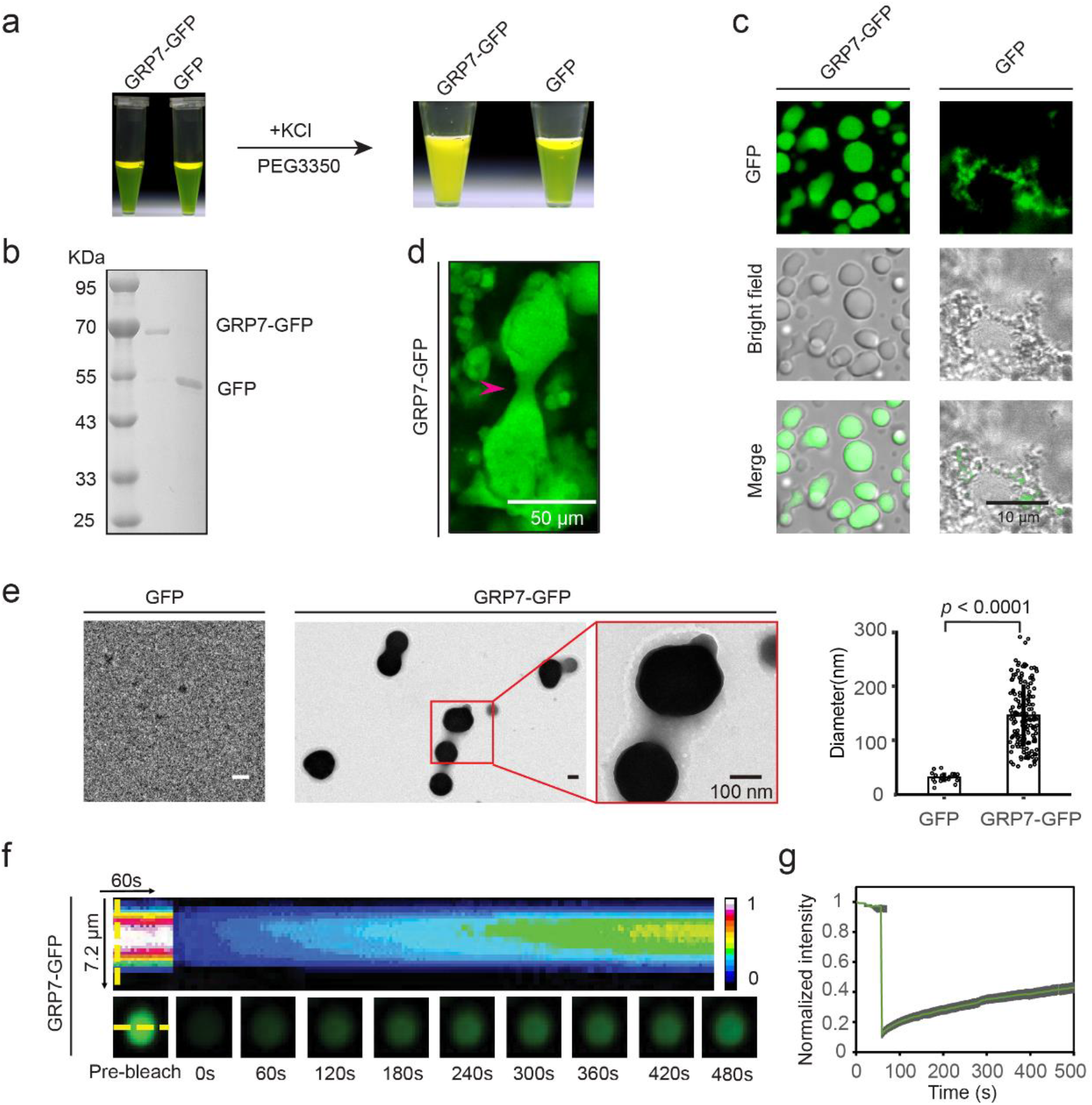
GRP7 undergoes phase separation *in vitro*. **a**, Solutions of 50 μM purified GRP7-GFP and GFP. Upon addition of 10% (w/v) PEG3350 and 100 mM KCl, the GRP7-GFP solution becomes turbid. **b**, Coomassie-stained SDS-PAGE gel of purified recombinant GRP7-GFP (∼63 kDa) and GFP (∼45 kDa). **c**, GRP7-GFP forms liquid-like droplets in an *in vitro* phase-separation assay. GRP7-GFP and GFP were at a concentration 50 μM with 100 mM KCl and 10% (w/v) PEG3350 added. **d**, Gel-like aggregates displayed fusion behaviours (magenta arrowhead). GRP7-GFP (70 μM) was mixed with 100 mM KCl and 10% (w/v) PEG3350 (see also Movie 4). **e**, Gel-like GRP7-GFP droplet (middle) imaged by transmission electron microscopy (TEM). The concentration of GFP was the same as that in the control (left). The sizes of GRP7-GFP and GFP condensates are shown at right. *P* values were calculated by one-way ANOVA. **f**, Representative kymograph heatmap showing the fluorescence intensity within a bleached GRP7-GFP droplet *in vitro* over time. **g**, Fluorescence intensity after FRAP of GRP7-GFP droplets *in vitro*. Data are shown as means ± s.d. values (n = 11).

To determine whether GRP7 itself can drive of its LLPS *in vivo*, we heterologously produced and purified recombinant GRP7 tagged with GFP in *E. coli*, using recombinant GFP alone as negative control (see Methods) (Fig. 3a,b). We then screened for conditions suitable for GRP7-GFP LLPS using a high-throughput phase separation screening (HiPSS) kit (Union-Biotech, Shanghai; see Methods). We observed the formation of GRP7-GFP spherical condensates under many conditions (for representative images, see Supplementary Fig. 4). We selected solutions containing PEG3350 and NaCl or KCl to use in characterizing GRP7-GFP LLPS *in vitro*. In solutions containing 100 mM KCl and 10% PEG3350, GRP7-GFP formed spherical, gel-like green puncta, whereas similar concentrations of GFP formed amorphous precipitates (Fig. 3c). Gel-like droplets fused upon contact (Fig. 3d, Movie 4) and small droplets coalesced into increasingly larger structures (Fig. 3d, Movie 5) in a process that was sensitive to crowding agents and pH (Supplementary Fig. 5a,c,d). Furthermore, 1,6-hexanediol disrupted LLPS of GRP7-GFP, suggesting that aromatic residues in the glycine-rich domain contribute to LLPS (Supplementary Fig. 5b, ref.^28^). To detect the nanoscale structure of GRP7-GFP droplets, we observed phase-separated specimens by transmission electron microscope (TEM). The droplets averaged ∼146 nm in diameter and had two layers: an even, dense inner layer and an outer layer that contained holes and was thin and sponge-like (Fig. 3e). No GFP-only droplets developed under the same conditions (Fig. 3e). TEM also revealed the fusion of two or more droplets (Fig. 3e, magnified image), consistent with the above microscopy results. Furthermore, FRAP analysis indicated that GRP7-GFP molecules diffused within droplets and were exchanged between droplets and the surrounding solution (Fig. 3f,g).

**Fig. 4.**
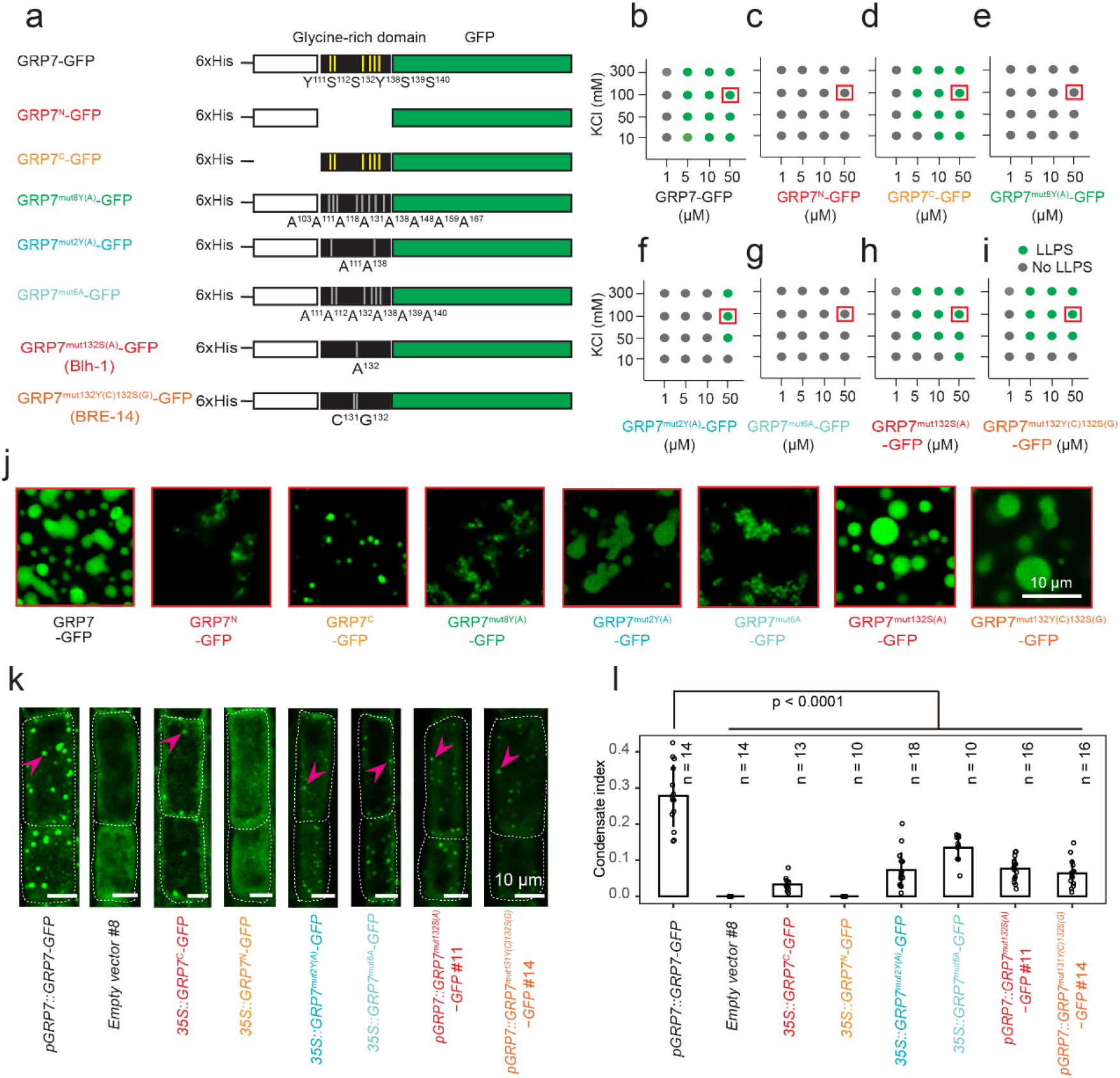
FER-dependent phosphorylation mediates GRP7 phase separation. **a**, Schematic representation of the GRP7 truncations and point mutations used in this study. Yellow bars highlight the FER-phosphorylation sites highlighted by Wang et al (2020). Gray bars denote mutated sites in this study. **b**-**g**, Phase-separation diagrams of full-length GRP7 (**b**), truncated GRP7 (**c, d**) and phosphorylation site mutant versions of GRP7 (**e**-**i**). **j**, Representative images of full-length, truncated and phosphorylation site mutant version of GRP7-GFP under the indicated conditions marked with red squares in **b**-**g**. Representative data from three independent experiments. **k**, GRP7 condensates in the cytoplasm of root cells from 5-d-old transgenic seedlings treated with NaAsO_2_ for 1 h. Magenta arrowheads point to GRP7 condensates. **l**. Condensate index of transgenic plants in **k** shown as means ± s.d. *P* values were calculated by one-way ANOVA.

**Figure 5.**
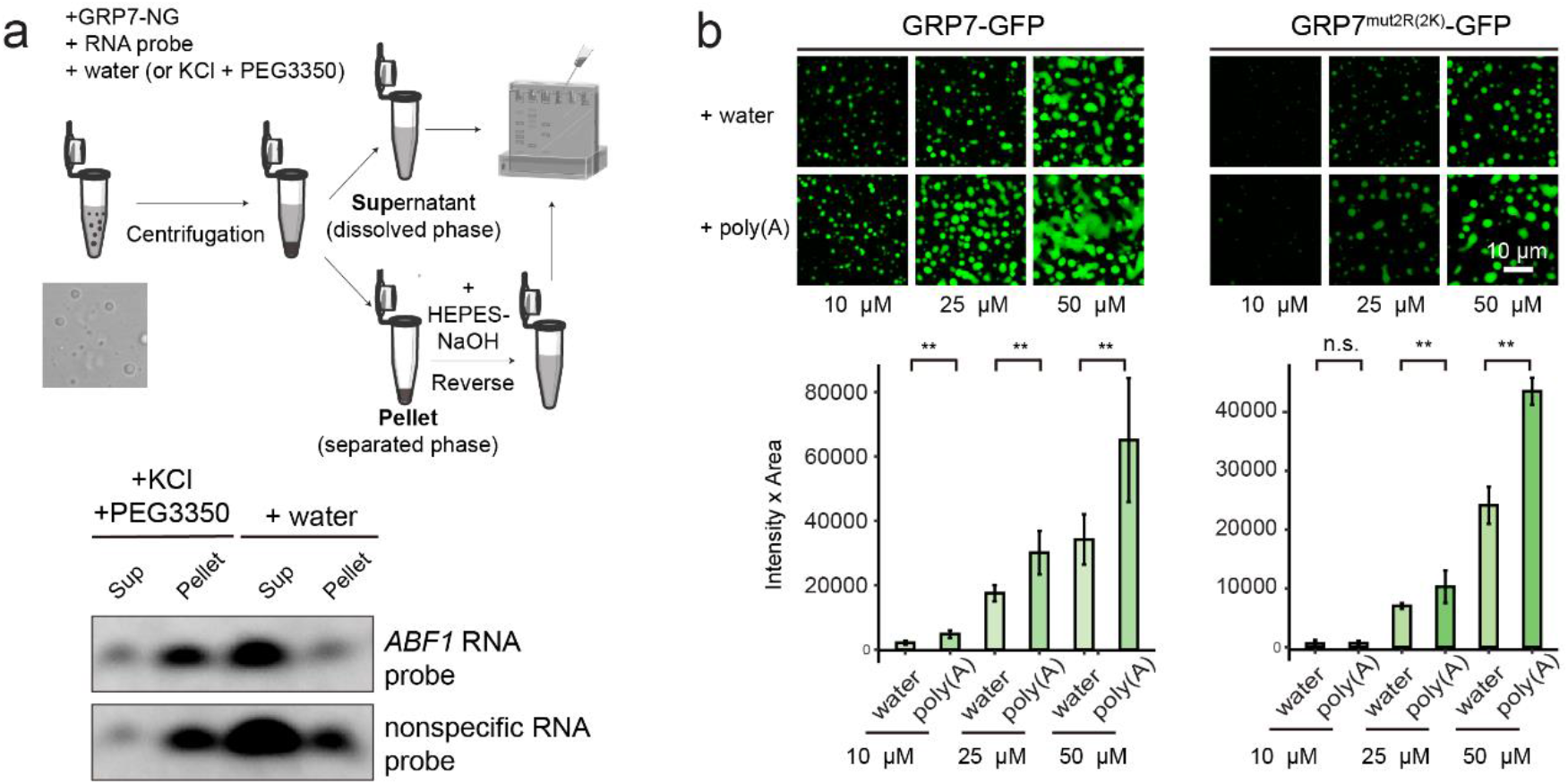
GRP7 phase separation facilitates RNA assembly. **a**, Phase-separated GRP7-NG droplets recruit RNA probes. A FITC-labelled probe (0.2 pM) was incubated with GRP7-NG protein (100 µL) for 10 min, together with water (control) or 100 mM KCl and 10% (w/v) PEG3350 added to form a phase-separated solution. The pellet was resuspended in 50 µM HEPES-NaOH, pH 7.5. Supernatant and pellet-reversed solution were loaded on a native gel (below). **b**. Poly(A) RNA promotes phase separation of GRP7-GFP and GRP7^mut2R(2K)^-GFP. Sums of the mean fluorescence intensity × area for all droplets per field of view from three images are shown (below). Data are shown as means ± s.d. values; \*\**P* < 0.01, n.s., nonsignificant by ANOVA (n=3).

To identify the domains of GRP7 responsible for LLPS, we generated a truncated GRP7 protein fused with GFP at its N-terminus (GRP7^N^, amino acids [aa] 1-86) and a GFP-tagged glycine-rich domain (GRP7^C^, aa 87-175) (Fig. 4a). As the phase-separation capacity of GRP7-GFP was strongly dependent on its own concentration and the solution ion concentration (Fig. 4a,b,j), we evaluated LLPS of these purified truncated GRP7s under the same conditions. Only the GFP-tagged glycine-rich domain (GRP7^C^), but not the N-terminal truncation (GRP7^N^), underwent LLPS, revealing that the glycine-rich domain is the main contributor to GRP7 LLPS (Fig. 4a,b,j). These results were consistent with the prediction of an IDR in the C-terminal glycine-rich region of GRP7 (Fig. 2a). Moreover, at the same concentration, GRP7-GFP droplets were larger than those formed by GRP7^C^-GFP (Fig. 4j), indicating that the N terminus of GRP7 also partially contributes to its phase separation. In line with the behaviours of GRP7-GFP phase separation *in vitro*, a transgenic line overexpressing *GRP7*^*C*^*-GFP* in the *grp7-1 8i* background displayed fewer, smaller condensates in root cells than *pGRP7::GRP7-GFP* transgenic lines, while we rarely detected condensates in the root cells of a line overexpressing *GRP7*^*N*^*-GFP* line in the *grp7-1 8i* background (Fig. 4k,l). Collectively, these data indicate that GRP7-GFP also undergoes LLPS both *in vivo* and *in vitro*.

### FER-dependent phosphorylation within GRP7 IDR is critical for GRP7 phase separation

FER can physically interact with and phosphorylate GRP7 at six sites in its glycine-rich domain^37^. Furthermore, treatment with RALF1, which activates FER kinase, facilitated GRP7-GFP condensate formation *in vivo* (Fig. 2b-d). We investigated whether FER-dependent phosphorylation of GRP7 influences its phase separation *in vitro* by generating three types of non-phosphorylatable GRP7-GFP variants and evaluating their behaviour in an *in vitro* phase-separation assay: GRP7^mut8Y(A)^-GFP (with eight Tyr-to-Ala mutations at the positions shown in Fig. 4a), GRP7^mut2Y(A)^-GFP (with two Tyr-to-Ala mutations were identified by Wang et al.^37^ as phosphorylated by FER and GRP7^mut6A^-GFP (with two Tyr-to-Ala and four Ser-to-Ala mutations at residues identified by Wang et al.^37^as phosphorylated by FER). Importantly, none of these non-phosphorylatable GRP7-GFP mutants phase separated as wild-type GRP7-GFP did (Fig. 4e-g), indicating that these phosphorylation sites are essential in regulating GRP7 phase separation. In line with the behaviour of GRP7-GFP phase separation *in vitro*, transgenic lines overexpressing *GRP7* ^*mut2Y(A)*^*-GFP* or *GRP7*^*mut6(A)*^*-GFP* in the *grp7-1 8i* background displayed weaker, smaller condensates in root cells (Fig. 4k,l). Collectively, these data indicated that FER-dependent phosphorylation may facilitate GRP7 phase separation.

We also generated and purified recombinant GRP7-GFP variants carrying the amino acid changes detected in the accessions Blh-1 (GRP7^mut132S(A)^-GFP) and BRE-14 (GRP7^mut131Y(C)132S(G)^-GFP) for evaluation in our *in vitro* phase-separation assay (Fig. 4a,h,j). Compared to intact GRP7-GFP, both GRP7^mut132S(A)^-GFP and GRP7^mut131Y(C)132S(G)^-GFP exhibited weaker phase-separation capacity (in the presence of 10 mM KCl and 10% PEG3350, Fig. 4b v.s. 4h-i). In line with this result, both the *pGRP7::GRP7*^*mut132S(A)*^*-GFP* #11 and *pGRP7::GRP7*^*mut131Y(C)132S(G)*^*-GFP* #14 transgenic lines had a lower condensate index than *pGRP7::GRP7-GFP* (Fig. 4k,l). These observations indicated that GRP7 variants from the accessions Blh-1 and BRE-14 have a reduced capacity to undergo phase separation, which may reflect their natural temperature habitat (Fig. 1c).

### GRP7 phase separation facilitates RNA assembly

To detect how GRP7 LLPS modulates plant response to temperature fluctuations, we explored whether GRP7 LLPS is involved in RNA assembly. Given that GRP7 functions in various RNA-processing events and contains an RNA-recognition motif, we hypothesized that liquid-like droplets of GRP7 might recruit RNA. Accordingly, we evaluated whether phase-separated GRP7 could recruit the *ABF1* (*ABSCISIC ACID RESPONSIVE ELEMENT-BINDING FACTOR1*) mRNA (encoding a basic leucine zipper transcription factor; ref.^39^), previously shown to bind to GRP7^37^. We mixed a synthetic *ABF1* RNA probe or a nonspecific RNA probe (which could not bind to GRP7; data not shown) with GRP7-NG (GRP7 not fused with GFP, to prevent any interference of GFP fluorescence with the RNA probe signal), PEG3350 and KCl. After centrifugation, we reversed the phase separation of the pelleted material was by resuspension in HEPES-NaOH buffer (50 μM, pH 7.5) before loading aliquots from the pellet and supernatant onto a nondenaturing gel (Fig. 5a, top). We detected enrichment of both the specific and nonspecific RNA probes in the pellet derived from phase separation; by contrast, in the absence of LLPS (with water added instead of PEG3350 and KCl), we observed a strong signal for both probes in the supernatant (Fig. 5a, bottom). This finding indicated that GRP7 phase separation is involved in the sequestration of RNA.

To eliminate the contribution of individual GRP7 molecules to RNA binding, we compared the phase-separation features of intact GRP7-GFP and a variant with mutations in the RNA-binding domain (Arg47Lys and Arg49Lys [GRP7^mut2R(2K)^-GFP], which lacks RNA-binding activity; ref.^40^) when incubated with homotypic RNAs (Fig. 5b). Poly(A) (100 ng/μL, the same concentration used for the other homotypic RNAs) promoted phase separation of GRP7-GFP over a range of 10 μM to 50 μM of protein and poly(C) did so at 10 μM and 50 μM, whereas poly(G) inhibited phase separation and neither poly(U) nor transfer RNA (tRNA) showed any obvious effect on phase separation (Fig. 5b, Supplementary Fig. 6). We selected poly(A) for evaluation GRP7^mut2R(2K)^-GFP. As with GRP7-GFP, poly(A) promoted phase separation of GRP7^mut2R(2K)^-GFP over a range of 25-50 μM (Fig. 5b) indicating that phase-separated GRP7 can sequester RNA, although less efficiently, even though it lacks the RNA binding capacity of individual molecules. Moreover, these results also suggested a specificity for GRP7 phase separation dependent on binding to RNA that contains poly(A) sequences, such as mRNA. Collectively, these results indicate that GPR7 condensates recruit RNA *in vitro*.

### GRP7 phase separation recruits two cold-shock proteins and a component of the translation machinery, reducing translational output

To uncover the molecular function of GRP7 condensates, we immunoprecipitated GRP7-GFP with an anti-GFP antibody from *GRP7-GFP* seedlings mock-treated or treated with NaAsO_2_ to induce LLPS, and then performed quantitative liquid chromatography (LC)-mass spectrometry (MS) analysis of co-purified proteins (Table S1). From samples with good reproducibility, we detected 47 proteins in NaAsO_2_-treated seedlings but not in mock-treated controls. Gene Ontology (GO) term analysis revealed an enrichment for stress-associated proteins in the Biological Processes (Supplementary Fig. 7a,b). CSP1 was among the proteins detected in GRP7-GFP cytoplasmic condensates, whose abundance rose upon cold exposure, which was correlated with improved translation of ribosomal protein mRNAs^12^. We hypothesized that GRP7-containing SGs may sequester CSP1 during cold stress. To test this idea, we performed split luciferase (split-LUC) complementation assays with GRP7 and CSP1 and its closest homolog, CSP3, finding that both CSP1 and CSP3 interacted with GRP7 (Fig. 6a, Supplementary Fig. 8a). We then evaluated *in vitro* GRP7 phase separation in the presence of CSP1 (Fig. 6b,c,e, Supplementary Fig. 9) or CSP3 (Supplementary Figs. 8b and 9b). Indeed, GRP7-GFP droplets incorporated full-length CSP1 and CSP3 fusion proteins labelled with mCherry (Fig. 6b,c, Supplementary Fig. 8b and 9) and also recruited free CSP1 and CSP3 molecules. (Fig. 6e, Supplementary Fig. 8c). These results showed that phase-separated GRP7 incorporates the mRNA chaperones CSP1 and CSP3.

**Fig. 6.**
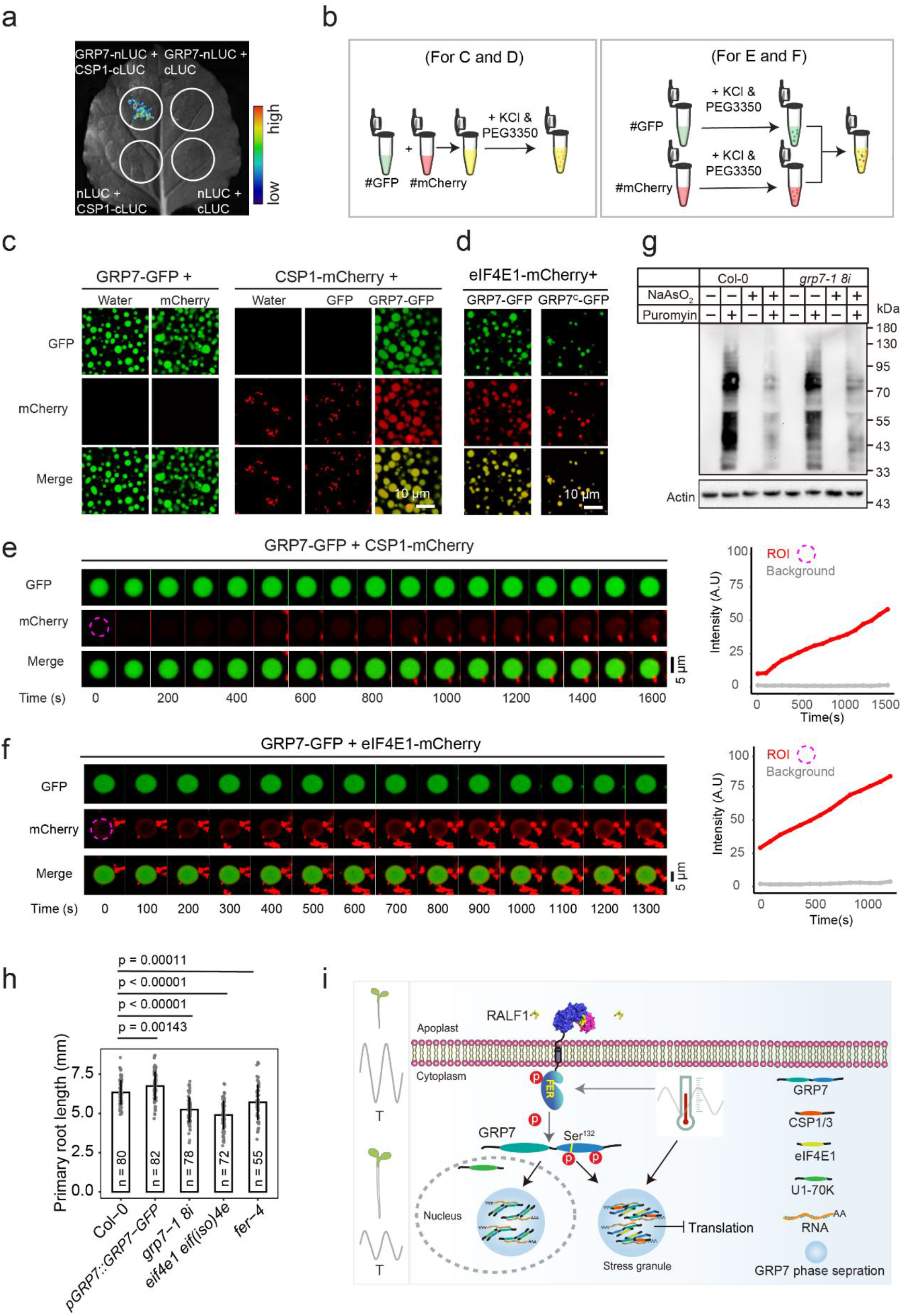
GRP7 endows seedlings with temperature resilience via the LLPS-mediated recruitment of translation machinery components. **a**, Split luciferase complementation assays showing the interaction between GRP7 and CSP1. The indicated constructs were co-infiltrated to *N. benthamiana* leaves via Agrobacterium-mediated infiltration. Luciferase activity was determined at 48 h after infiltration. **b**, Schematic diagram illustrating sample preparation for the experiments shown in **c**-**f. c**,d, GRP7-GFP droplets recruit CSP1-mCherry (**c**) and eIF4E1-mCherry (**d**). Liquid droplets formed after mixing 50 µM CSP1 -mCherry with GRP7-GFP and incubating for 10 min at room temperature. **e, f**, Phase-separated GRP7-GFP droplets recruit the free forms of CSP1-mCherry (**e**) and eIF4E1-mCherry (**f**). A phase-separated GRP7-GFP solution was mixed with a CSP1-mCherry or eIF4E1-mCherry solutions, and images were acquired immediately. Regions of interest are highlighted by the dashed magenta circles. The graphs at right in each panels show changes in the intensity of CSP1-mCherry and eIF4E1-mCherry over time. **g**, Protein synthesis in Col-0 and *grp7-1 8i* seedlings treated with NaAsO_2_. Five-day-old seedlings were treated with liquid half-strength MS medium containing 500 µM of NaAsO _2_ for 1 h and then incubated with the same medium containing 10 mg/mL puromycin for 4h. Total proteins were tested by immunoblotting. **h**, Primary root length of Col-0, *pGRP7::GRP7-GFP, grp7-1 8i, eif4e1 eif(iso)4e*, and *fer-4* seedlings grown under fluctuating temperatures (as in Fig.1d) for 5 d. Data are shown as means ± s.d.. *P* values were calculated by one-way ANOVA. **i**, Model for the mechanism of plant adaptation to temperature fluctuation. The RALF1-activated receptor kinase FER phosphorylates GRP7 at Ser132 in its glycine-rich domain, which facilitates LLPS of GRP7 to promote the assembly of RNA and of eIF4E1, CSP1 and CSP3 in the cytoplasm, leading to an inhibition of translation to control root growth. The FER-GRP7 LLPS module can be modulated by fluctuating temperatures.

As GRP7 forms puncta in the cytosol (Fig. 2b-d), we speculated that the GRP7 condensates may incorporate certain components of the translation machinery as well. To test this hypothesis, we evaluated the GRP7-interacting protein eukaryotic translation initiation factor 4E1 (eIF4E1), a key component of translation machinery^41^, in two *in vitro* GRP7 phase separation assays (Fig. 6d and Supplementary Fig. 10a). Indeed, GRP7-GFP droplets incorporated full-length eIF4E1 (Fig. 6d and Supplementary Fig. 10a). In the FRAP assays, GRP7-GFP mixed with eIF4E1-mCherry recovered from photobleaching more slowly than GRP7-GFP alone (Supplementary Fig. 10b), revealing that addition of eIF4E1 altered GRP7-GFP molecular dynamics. Moreover, GRP7-GFP mixed with eIF4E1-mCherry at a concentration ratio of 2.5:1 formed larger droplets than similar mixtures at different ratios (Supplementary Fig. 10a). Although eIF4E1-mCherry did not phase separate under such conditions, its free molecules were recruited into GRP7-GFP droplets (Fig. 6f). These results showed that phase-separated GRP7 incorporates a component of the translation machinery (eIF4E1) and RNA chaperones (CSP1 and CSP3) in the cytoplasm, which may drive mRNA translation. Similarly, phase-separated GRP7 functions in incorporated a component of the spliceosome (U1-70K) in the nucleoplasm (see Supplementary Text. Supplementary Figs. 11 and 12).

To test if translation was blocked by the formation of GRP7 condensates, we investigated total protein synthesis in Col-0 and *grp7-1 8i* that were treated with NaAsO_2_ (to rapidly induce GRP7 condensate formation). The protein synthesis rate of *grp7-1 8i* seedlings was lower than that of Col-0 under normal conditions, but rose above that of Col-0 upon treatment with NaAsO_2_ (Fig. 6g). This result indicated that GRP7 condensate can sequester eIF4E1 and/or CSP1 and CSP3 to inhibit translation under stress conditions. Correspondingly, a double mutant lacking eIF4E1 function, *eif4e1 eif(iso)4e* ^42^, displayed a root length comparable to those of the of *grp7-1 8i* and *fer-4* mutants in a cold response assay, all of which were shorter than that of Col-0 seedlings (Fig. 6h). Taken together, these results indicate that the FER-GRP7 module endows plants with enhanced temperature resilience via LLPS-mediated recruitment of translation-related components.

## Discussion

Although GRP7 participates in various steps of RNA processing^37, 43-45^, flowering time control^43, 46^, the immune response^41^, and the circadian clock^47, 48^, its mechanism involved in temperature responses remains unclear. Here, we provide experimental evidence that LLPS plays a key role in GRP7-dependent response to temperature fluctuations: 1) GRP7 undergoes LLPS and forms thermally inducible SGs *in vivo*; 2) the C-terminal glycine-rich domain of GRP7 is the main contributor to its LLPS; 3) within the IDR, a FER-dependent phosphorylation site, which is also subject to natural variation, determines GRP7 phase separation behaviours, as the Col-0 GRP7 variant has a higher capacity for phase separation that correlates with plant adaptation to a wider temperature range; 4) GRP7 condensates incorporate RNA, the translation machinery component eIF4E1 and the two RNA chaperones CSP1 and CSP3 to inhibit translation (Fig. 6i). Conceptually, therefore, GRP7 might act as *scaffold* and RNA, eIF4E1, CSP1 and CSP3 as *clients* in the condensates, according to the scaffold-client model^49^. Our findings improve the understanding of how GRP7 confers greater adaptation to a wider temperature range and unveiled a synergistic relationship between GRPs and CSPs.

Phase separation has been implicated in the regulation of numerous processes in plants, including transcription (FLOWERING CONTROL LOCUS A [FCA], ref.^50^), microRNA processing (SERRATE [SE], ref.^51^), phytohormone sensing (Auxin Response Factor 7 and 9 [ARF7 and ARF19], ref.^19^), adaptation to higher temperatures (EARLY FLOWERING3 [ELF3], ref.^17^), cargo sorting (SORTING OF cpTat SUBSTRATES TO THYLAKOID MEMBRANES1, [STT1 and STT2], ref.^52^), pathogenic responses (NONEXPRESSOR OF PATHOGENESIS-RELATED GENES1 [NPR1], ref.^21^; guanylate-binding protein-like GTPase 1 and 3 [GBPL1 and GBPL3], ref.^20^), autophagy (AUTOPHAGY-RELATED 8e [ATG8e], ref.^53^), the circadian clock (Cryptochrome 2 [CRY2], ref.^54^), timing of meiotic transition (MEIOSIS ARRESTED AT LEPTOTENE2 [MEL2], ref.^55^) and seed germination (FLOE1, ref.^18^, reviewed in ref.^30, 56^). The key driver proteins exhibiting phase separation are RNA-binding proteins (RBPs) (FCA, MEL2), zinc-finger proteins (SE,), transcription factors (ARF7 and ARF19, NPR1) or transcriptional regulators (ELF3), autophagosome proteins (ATG8e), GTPases (GBPL1 and GBPL3), photoreceptors (CRY2) and ankyrin-repeat proteins (STT1 and STT2). An increasing number of RBPs containing typical IDRs have been shown in animals to undergo phase separation. These proteins play essential roles in diverse physiological and pathological processes^14, 15^. The mechanism by which their LLPS-related biochemical behaviours are regulated and their relationships with stress or disease states are coming into focus. To date, only two plant RBP, FCA and MEL2^30, 50, 55, 56^, have been documented to display LLPS behaviours; these function in precursor mRNA polyadenylation in Arabidopsis^50^and that regulates the timing of meiotic transition in rice (*Oryza sativa*)^55^, respectively. LLPS of GRP7, reported here, provides a novel example of a plant RBP that mediates signal transduction in response to temperature stress.

Little is known about the role of the glycine-rich domain, which exhibits the characteristics of an IDR. *In vitro* RNA-binding assays have shown that truncations of this domain reduce the RNA-binding activity of the protein^13^. We found that (1) the glycine-rich domain drives phase separation of GRP7 to recruit other partner proteins, including CSP1, CSP3, eIF4E1 (Fig. 6) and U1-70K (see Supplementary Text, Supplementary Figs. 11 and 12); (2) phosphorylation sites within this domain are phosphorylated by a receptor kinase; and (3) the glycine-rich domain can bind RNA on its own, even after the GRP7 RNA-binding site is deleted. These results suggest a novel role for the glycine-rich domain of GRP7. Future studies will be necessary to determine the interactions between the RNA-binding and the glycine-rich domain.

Temperature fluctuations inhibits root cell growth, causing a drop in crop productivity. Rapid, effective restoration after short-term extreme cold or heat stress is essential for plant growth. The RALF1-FER module has been widely considered a key node to steer cell growth and is involved in multiple cross-talking signalling pathways, including temperature^35, 57^. FER was recently reported to harbor a temperature-sensitive site (Gly 41) in its extracellular domain, which does not affect its perception to external signals at normal temperatures (20 °C), but abolishes the formation of root hairs at elevated temperatures (30 °C, ref.^36^). FER is required for perceiving limited nutrients availability caused by low temperature, to interacts with and activate TORC1-downstream components to trigger root hair growth^33^. Future work will be needed to precisely define whether thermally inducible GRP7 phase separation is directly under FER control. Due to its multiple functions, FER has recently emerged as an integrator for nutrient signalling^33^ and a potential target for crop improvement and protection^57^. Our previous^37^ and current studies suggest that FER aids Arabidopsis root growth after sharp temperature changes by regulating of GRP7 LLPS. This mechanism offers a potential strategy whereby LLPS of RBPs enables plants cells to respond rapidly to temperature variation. Chemicals able to induce GRP7 condensates may pave the way to modulating of temperature resilience in crops.

## Methods

### Plant growth condition

Arabidopsis (*Arabidopsis thaliana*) seeds were first surface-sterilized with 15% (v/v) NaClO for 5 min and then stratified for 2 d at 4 °C in darkness to break dormancy. Seeds were sown on solidified half-strength Murashige and Skoog (MS) medium (Murashige and Skoog basal salt mixture [PhytoTechnologies Laboratories], pH of 5.7, containing 0.8% [w/v] agar [Difco] and 1% [w/v] sucrose [Sigma-Aldrich]). Unless indicated otherwise, seedlings were grown at a constant temperature of 22 °C with long-day (16-h light/8-hdark) photoperiod. When seedlings were exposed to fluctuating temperatures, plates were placed into a growth chamber set to various temperatures from 10 °C to 34 °C (interval = 2 °C per h) for 5 d under a similar photoperiod (see Fig. 1d for temperature program). To calculate primary root length, at least 40 seedlings were measured with FIJI software.

### Plasmid construction for heterologous protein production

First, the coding sequence of *GRP7* fragment or its mutated variants was inserted into the pCAMBIA2300-GFP plasmid using restriction sites *Xba*I and *Bam*HI. Second, the coding sequence of *GRP7* or its mutated variants plus the GFP tag connected by a short linker (7 amino acids [aa]) or *GFP* alone were cloned into the *Eco*RI and *Bam*HI restriction site of the pET32a expression vector (Novagen) containing a thioredoxin (Trx) tag (109 aa, to improve the solubility of the recombinant protein), S tag and His tag. Except for minor modifications described below, eIF4E1-, U1-70K-, CSP1- and CSP3-related constructs were built and the corresponding recombinant proteins were purified using similar methods described above for GRP7. The coding sequences for *CSP1, CSP3, eIF4E1* and truncated *U1-70K* (589 bp to 1,281 bp) fragments were cloned in frame with the mCherry sequence using bridging PCR. The resulting mCherry fusion cassettes were cloned into the *Eco* I and *Bam*HI restriction sites of pRSF-Duet (His tag) and pGEX-4T-1 (GST tag; note: U1-70K-mCherry with His-tag could not be enriched with Ni-NTA beads for an unknown reason.)

### Plasmid construction for plant experiments

For *Arabidopsis* transgenic lines, the coding sequences of *GRP7* variants (*GRP7*^*mut132S(A)*^, *GRP7*^*mut131Y(C)132S(G)*^, *GRP7*^*mut2Y(A)*^) variants were synthesized by Qsingke Company. The coding sequences of *GRP7*^*C*^, *GRP7*^*N*^, and *GRP7*^*mut2Y(A)*^ were cloned into the pCambia-2300 vector. *GRP7*^*mut132S(A)*^, *GRP7mut*^*131Y(C)132S(G)*^ and *GRP7*^*mut2Y(A)*^ were cloned in frame and upstream of the *GFP* sequence and placed under the control of the *GRP7* promoter in the pCambia-1300.

For split luciferase complementation assays, the coding sequence of *GRP7* was inserted into pCAMBIA1300-NLuc (pNL), while the coding sequences of *CSP1* and *CSP3* were cloned into pCAMBIA1300-CLuc (pCL), using restriction sites XbaI and BamHI. The resulting constructs were named pCambia1300-CSP1-cLUC, pCambia1300-CSP3-cLUC and pCambia1300-GRP7-nLUC.

For root protoplasts transformation and transient expression in *Nicotiana benthamiana* leaves, the fusion cassettes *eIF4E1-mCherry* and *DCP1-mCherry* were inserted into the pHB vector using restriction sites *Xba*I and *BamH*I.

### Arabidopsis transgenic plants

Transgenic plants were generated by Agrobacterium (*Agrobacterium tumefaciens*)-mediated (GV3101 strain) transformation of *grp7-1 8i* (*grp7-1* mutant stably expressing an RNAi construct targeting *GRP8*). Primary transformants (T_1_) were selected based on hygromycin resistance, and T_2_ lines containing only one T-DNA insertion were selected for further characterization by determining the Mendelian segregation ratio (3:1) of hygromycin-resistant seedlings in the T_2_ progeny and confirmed in the T_3_ progeny. The GRP7^mut6A^-GFP line was described in our previous work^37^.

### Recombinant protein production in *Escherichia coli*

The appropriate constructs listed above were transformed into *Escherichia coli* (*E. coli*) BL21 star (DE3) cells. Single clones were picked and transferred to LB medium for overnight culture at 37 °C with shaking at 250 rpm. The cultures were diluted 1:50 into fresh LB medium containing 50 mg mL^-1^ ampicillin (for pGEX-4T-1 and pET32a) or kanamycin (for pRSF-Duet) for growth until reaching an optical density at 600nm (OD_600_) of 0.8. Protein production was induced with 0.5 mM isopropyl-β-D-thiogalactopyranoside (IPTG) at 16 °C overnight. The cell cultures were spun down at 5,000 rpm, and the cell pellet was resuspended in lysis buffer (25 mM Tris-HCl, pH 7.5, 300 mM NaCl, 20 mM imidazole, 1 mM phenylmethylsulfonyl fluoride [PMSF]) and lysed with an ultrasonic cell disruptor on ice. The lysates were centrifuged at 12,000 rpm for 10 min at 4 °C, and the resulting supernatant was transferred to a new tube and incubated with Ni-NTA resin at 4 °C for 3 h. The Ni-NTA beads were washed extensively with wash buffer (20 mM Tris-HCl, pH 7.5, 300 mM NaCl, 30 mM imidazole), and recombinant proteins were eluted with elution buffer (20 mM Tris-HCl, pH 7.5, 300 mM NaCl, 300 mM imidazole). The eluates were desalted via centrifugation with a salt-free buffer (50 mM HEPES-NaOH (pH 7.5)) using Amicon Ultra centrifugal filters (30K MWCO, Millipore). The centrifugation step was repeated five times, to achieve removal of 99.9% of salts. The protein concentration was measured using a BCA kit (PC0020-500, Solarbio) according to the user manual and finally adjusted to 100 μM. Aliquots of 100 μM GRP7 proteins were then stored in 50 mM HEPES-NaOH, pH 7.5, at −80 °C until use.

### Visualization of phase separation *in vitro* and *in vivo*

The high-throughput phase-separation screening (HiPSS) kit comprised 184 conditions, including a variety of electrolytes and polymers to simulate a crowded cell environment, as well as numerous pH values and charged reagents (for information on all components in each well, see http://en.union-biotech.com/product/4335839/). This kit was used to screen proteins with phase-separation capacity, evaluate the strength of their phase separation and identify other molecules (such as proteins, nucleic acids or small molecules) that influence the phase separation of proteins of interest. For *in vivo* visualization, seedlings were grown vertically on half-strength MS plates supplemented with 1% (w/v) sucrose and 0.5% (w/v) Phytagel (Sigma-Aldrich, P8169). Five-day-old seedlings were moved to liquid half-strength MS medium containing 2 μM synthetic RALF1 peptide (Sangon Biotech Co. Ltd). After the indicated times of treatment, subcellular localization analyses were performed with a Zeiss LSM880 confocal microscope using a 63× oil immersion objective and the GaAsP spectral detector of the LSM880. GFP was excited at 488 nm and detected at 491-535 nm. All images are z-stack maximum projections using a step size of 0.45 μm, spanning the entire width of the nucleus. For *in vitro* visualization, the protein solution was loaded into a glass-bottom cell culture dish. The dishes were then imaged with a Zeiss LSM880 confocal microscope as described above.

### Condensate index measurements

Images for *in vivo* GRP7 condensate estimation were analysed with FIJI software to calculate condensate index (CI). Epidermal cells in the transition zone of the root were imaged. The condensate size and density were estimated by calculating CI, defined as: CI = N*As*C/A/Bi, where N is the number of condensates per cell; As and A are the aera of the selected cell and the average size of GRP7 condensates, respectively; and C and Bi are the average fluorescence intensity of GRP7 condensates and of the selected cell, respectively.

### Fluorescence recovery after photobleaching (FRAP)

*In vivo* and *in vitro* FRAP experiments were carried out with a Zeiss LSM 880 microscope. Droplets were bleached with a 488-nm laser pulse (three repeats, 100% intensity). Recovery from photobleaching was recorded for the indicated times. Image analysis was performed with FIJI.

### Prediction of intrinsically disordered regions (IDRs)

IDR and PrLD predictions for GRP7 were performed using PONDRE (http://www.pondr.com) and PLAAC (http://plaac.wi.mit.edu/) with full-length protein sequences.

### Transmission electron microscopy

GRP7-GFP (50 μM) was used in the *in vitro* phase-separation assay. Phase-separated GRP7-GFP (10 μL) was mounted onto carbon-coated grids. Then, the grids were then air-dried at room temperature for 1 min and rinsed three times with distilled water. The excess liquid was wicked away with dust-free filter paper. Then, 10 μL of 1% (w/v) phosphotungstic acid was spotted on the carbon-coated grids, and dyed for 30-60s. Dust-free filter paper was used to wick away excess liquid, and then the samples were then dried and observed under a microscope. Images were acquired with a JEM-2100Plus transmission electron microscope.

### RNA recruitment assays

Fluorescein isothiocyanate (FITC)-labelled RNA probes were supplied by TsingKe Biological Technology Co. Ltd. The sequences of the ABF1 and nonspecific RNA probe were 5’-CUUACUCCGUGCUGGCGUUGUUAAAGAAGA-3’ and 5’-AACCCUUUCACGGAGGAAGAAGAAGAAAGGCUUUUAGC-3’, respectively. To perform phase separation, the FITC-labelled probe (0.2 pM) and 50 μM of purified recombinant GRP7-NG protein were added to mixtures of PEG3350 (10%, w/v) (or the same volume of water as the negative control) and KCl (100 mM). The solutions were incubated for 10 mins and were then centrifuged at 12,000 rpm for 5 mins, at room temperature. The supernatants were transferred to new tubes, and the pellets were diluted into a volume equal to that of the supernatant with HEPES buffer (pH 7.5). To prevent RNA degradation, 1 μL of a ribonuclease (RNase) inhibitor (Thermo Fisher, N8080119) was added. Next, equal volumes of supernatant and the diluted pellet mixture were loaded onto a 4% (w/v) polyacrylamide and 0.5× Tris-borate EDTA native gel, which was run for 40 min and then exposed in a fluorescence imager plate reader.

### Root growth inhibition assays

For the RALF1 root growth inhibition assay, 3-d-old seedlings with comparable growth vigour were transferred to liquid half-strength MS medium containing 4 μM RALF1 for 2 d, and their root length was then analysed as previously described^37^.

### Split luciferase complementation assay

Agrobacterium strain GV3101 harbouring pCAMBIA1300-CSP1-cLUC, pCAMBIA1300-CSP3-cLUC and pCAMBIA1300-GRP7-nLUC constructs were incubated in LB liquid medium containing 150 μg mL^-1^ rifampicin and 50 μg mL^-1^ kanamycin for 24 h. The bacteria were collected by centrifugation at 4,000 rpm for 5 min and the pellets were resuspended in infiltration buffer (10 mM MgCl_2_, 10 mM MES pH 5.6, and 100 μM acetosyringone) at a final concentration of OD_600_ = 1.0. The cell suspensions were kept at room temperature for 3 h before infiltration. For co-infiltration, equal volumes of two different strains carrying the indicated *nLUC* and *cLUC* constructs were mixed prior to infiltration. The bacteria were infiltrated into young leaves of *Nicotiana. benthamiana* by using a 1-mL disposable syringe and grown for 48 h-72 h before observation. The infiltrated leaves were painted with D-luciferin and imaged with a CCD camera (NEWTON 7.O Bio/FUSION FX–Plant Imaging Systems).

### Transient protoplast transfection

Root protoplasts were isolated from the roots of 7-d-old *35S::GFP* and *35S::GRP7-GFP* seedlings by digestion with cellulase and macerozyme. The protoplasts were then transiently transfected with *eIF4E1-mCherry* or *DCP1-mCherry* construct and incubated for 16h in the dark. One hour before observation with microscope, the transfected protoplasts were treated with 500 μM NaAsO_2_ to induce LLPS.

### *N*.*benthamiana* leaf infiltration

The steps of bacterial cultivation and infiltration were as described above. For co-infiltration, equal volumes of two different strains carrying the indicated pCAMBIA2300-GRP7-GFP and pHB-eIF4E1-mCherry plasmids were mixed prior to infiltration. The fluorescence in the infiltrated leaves was detected with a Zeiss 880 microscope.

### Climate data

Climate data from the original collection sites of five Arabidopsis accessions (was extracted from either 1960-1990 average climate maps of Europe (monthly precipitation, minimum and maximum temperature^58^) or from a time series from 1958 to 2017 (monthly temperature^59^).

### Protein synthesis analysis

Five-day-old seedlings were treated with 500 μM of NaAsO_2_ for 1h. The seedlings were then incubated for 4 h in half-strength MS medium containing 10 mg/mL of puromycin. After treatment, seedlings were washed twice in liquid half-strength MS medium and harvested and ground to powder in liquid nitrogen. The lysate was mixed with SDS sample loading buffer and incubated at room temperature for 30 min. The mixed solution was centrifugated at 12,000 rpm for 10 mins. Equal amount in µg from each the supernatant was loaded on a 10% (w/v) SDS-PAGE gel and transferred to a PVDF membrane (Bio-Rad). Puromycin incorporation was measured using an anti-puromycin antibody (Merk Millipore).

### Cold shock test in *Escherichia coli*

BL21 cells transformed with *GFP* or *GRP7-GFP* plasmids were grown in LB medium containing ampicillin. One millilitre of the overnight cultures of the BL21 cells containing each construct were used to inoculate fresh LB medium (200 mL) containing 0.2 mM of IPTG and incubated at 16 °C. The growth rate was monitored by measuring the OD_600_.

### Statistics

Significant differences in data were analysed by two-tailed Student’s t tests or by multivariate comparisons (one-way analysis of variance, ANOVA) using R studio software. All statistical tests are clearly described in the figure legends and/or in the Methods.

## Supporting information

Combined supplemenatary data

## SUPPLEMENTAL INFORMATION

Supplemental Information can be found online at *Nature Chemical biology* Journal Online.

## AUTHOR CONTRIBUTIONS

F. Y. and F. X. designed the research; F. X., L. W., J. F. S., W.J. C., Y.B. L., and D. S. performed the experiments; F. X. and F. Y. wrote the manuscript.

## ACKNOWLEDGMENTS

The authors are grateful to Meixia Hu and Shufeng Song in State Key Laboratory of Hybrid Rice, Hunan Hybrid Rice Research Center for their technical support on confocal microscope. We thank Hui Chu in Institute of Chemical Biology and Nanomedicine, Hunan University for her their technical support on TEM. Jia Chen and other members in the laboratory are thanked for critical reading the manuscript and helpful discussions. Liang Wang in School of Life Sciences, Tsinghua University is thanked for helpful discussion.

