## Supplementary material for "The Receptor Kinase FER Mediates Phase Separation of Glycine-Rich RNA-Binding Protein 7 to Confer Temperature Resilience in *Arabidopsis*": Combined supplemenatary data

### Supplementary Materials

#### Supplementary Text

##### Cytoplasmic GRP7 condensates are stress granules

In this study, we concentrated on the GRP7 condensates in the cytoplasm. The pattern of cytosolic GRP7 condensates resembles that of stress granules (SGs, Figure 2 in ref.[60, 61]) and/or processing bodies (P-bodies, Figure 2 in ref.[61, 62]), which prompted us to determine whether GRP7 condensates are SGs or PBs. We first screened stress signals that can induce GRP7 condensation according to the published experimental conditions in both animals and plants. A shift from 22 °C to 38°C, or treatments with the pharmacological agents $NaAsO_2$ (500 µM for 1h), imidazole (500 µM for 2h), $CaCl_2$ (100 µM for 2h), PEG6000 (10% [w/v] for 1h) and $H_2O_2$ (1.0% [v/v] for 5h), induced cytoplasmic GRP7 condensates in root cells to various degrees (Supplementary Fig. 2a). We then induced GRP7 condensates in *Arabidopsis* seedlings and protoplasts expressing *GRP7-GFP* (for results see also Fig. 2) and *N. benthamiana* leaves transiently infiltrated with *GRP7-GFP* construct by treatment of RALF1 with $NaAsO_2$, a widely-used SGs chemical inducer in animal cells[15]. Not surprisingly, *GRP7-GFP* roots (Supplementary Fig. 3a) or protoplasts prepared from *GRP7-GFP* roots (Supplementary Fig. 3b) and treated with $NaAsO_2$ formed obvious GRP7 cytosolic condensates within 2 h. In a third step, we introduced a construct encoding the SG marker protein eIF4E1 (a eukaryotic translation initiation factor 4E1, ref.[61]), fused to mCherry, into the *GRP7-GFP* root protoplasts. After the induction of GRP7-GFP condensate, the GRP7-GFP signal colocalized with that of eIF4E1-mCherry (Supplementary Fig. 3c). Fourth, *N. benthamiana* leaves were infiltrated with a mixture of agrobacteria harbouring the *GRP7-GFP* and *eIF4E1-mCherry* constructs; condensates were induced by $NaAsO_2$ treatment. We observed the colocalization of GRP7-GFP and eIF4E1-mCherry for the cytosolic GRP7 condensates, whereas nucleoplasmic GRP7-GFP condensate did not uniformly overlap with the eIF4E1-mCherry signal (Supplementary Fig. 3d). These findings obtained from protoplasts and *N. benthamiana* leaves indicated that GRP7 LLPS forms SGs in the cytoplasm *in vivo*.

SGs serve as sorting sites, where mRNAs are targeted for storage, re-initiation or degradation by transfer to PBs[63]. Given that SGs and PBs have this functional continuity and that they are even physically connected in some cases[61], we also examined the relationship between GRP7 condensates (representing an example of SGs) and PBs. To this end, we introduced a contruct encoding the PB marker protein Decapping cofactor 1 (DCP1) fused with mCherry (DCP1-mCherry) into *GRP7-GFP* root protoplasts. Prior to $NaAsO_2$ treatment, GRP7-GFP almost uniformly localized to the outer layer of PBs, while GRP7-GFP fluorescence shifted to form puncta at the PB boundary after $NaAsO_2$ treatment (Supplementary Fig. 3e). This observation suggested that the formation of GRP7 condensate may correlated with mRNA processing or translation control through SGs and PBs.

**GRP7 phase separation also recruits U1-70K**

Given that alternative splicing also controls RNA metabolism and that GRP7 forms puncta in the nucleus (Fig. 2b-d), we speculated that the GRP7 condensates may incorporate certain components of the spliceosome. To test this hypothesis, the key components of the spliceosome (U1-70K), which were shown to interact with GRP7[37], were evaluated in two *in vitro* GRP7 phase-separation assays (Supplementary Fig. 11). Indeed, GRP7-GFP droplets incorporated full-length and truncated U1-70K labelled with mCherry (Supplementary Fig. 11c). In the FRAP assay, GRP7-GFP mixed with U1-70K$^{(197\text{-}427)}$-mCherry recovered from photobleaching faster than GRP7-GFP alone (Supplementary Fig. 11f), revealing that the molecular dynamics are altered by U1-70K. Phase-separated GRP7$^{C}$-GFP exhibited similar effects on the incorporation of U1-70K$^{(197\text{-}427)}$-mCherry (Supplementary Fig. 12). Since U1-70K$^{(197\text{-}427)}$-mCherry also phase-separates under the same conditions (10% [w/v] PEG3350 and 100 mM KCl) as

GRP7-GFP (Supplementary Fig. 11a,b), PEG3350 and KCl were added to these two proteins separately before they were mixed (Supplementary Fig. 11d right panel). GRP7-GFP droplets recruited U1-70K$^{(197\text{-}427)}$-mCherry droplets (Supplementary Fig. 11d). Together, these results indicated that phase-separated GRP7 also recruits U1-70K.

### Supplementary Figures

a

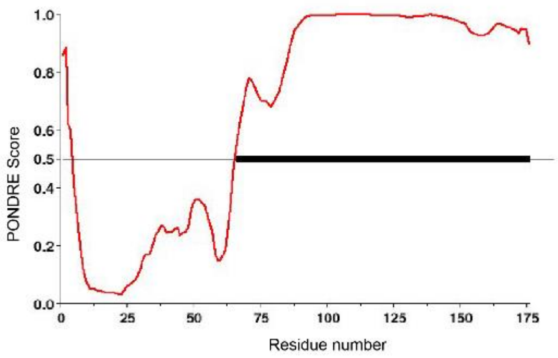


**Supplementary Fig. 1. Intrinsic disorder tendency of GRP7.**

**a**, Intrinsic disorder tendency of GRP7, as predicted by the PONDR online tool (http://www.pondr.com/). The most disordered region of the protein is represented as a black line.

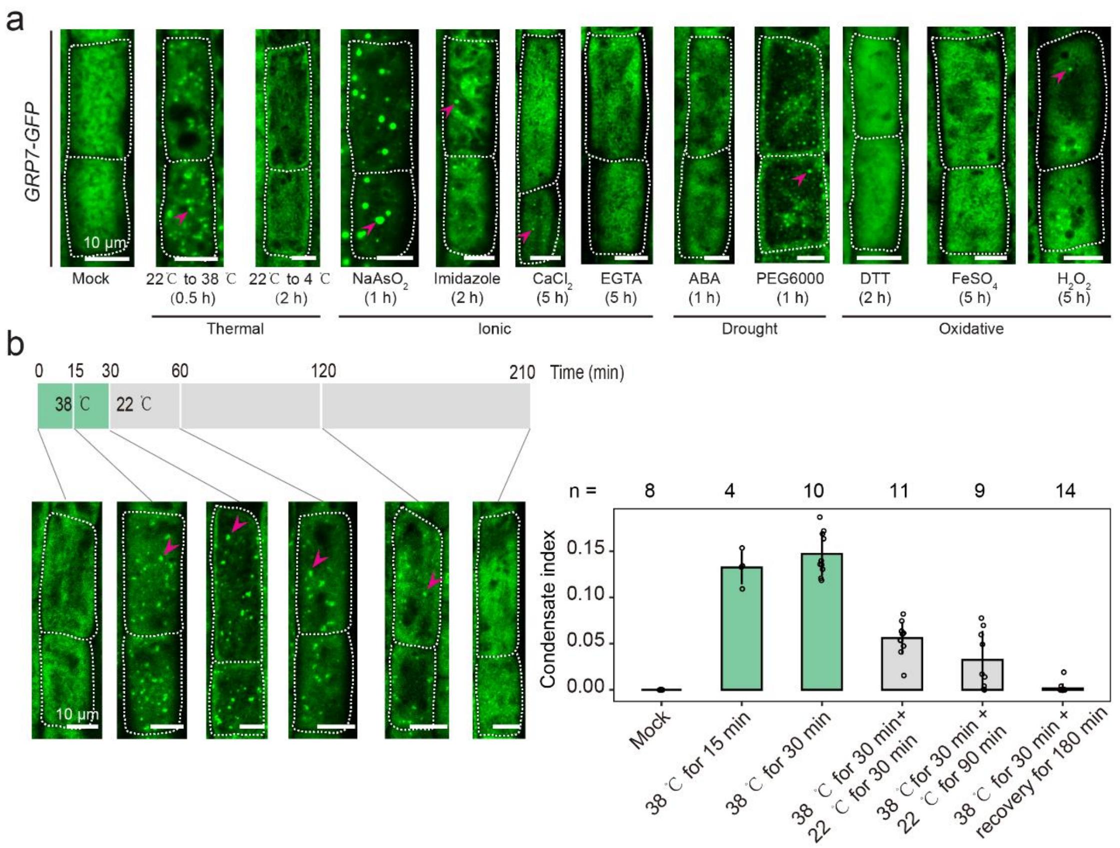


**Supplementary Fig. 2. Cytoplasmic GRP7 condensate induced by various conditions in root cells.**

**a**, Screening of external stress signals for their ability to induce GRP7 condensates in root cells. Five-day-old of GRP7-GFP seedlings were treated with temperature shift or with liquid half-strength MS medium containing the indicated chemicals for the indicated time periods. **b**, Heat-induced GRP7 condensate formation is reversible. Five-day-old of GRP7-GFP seedlings were exposed to warm temperature (38 °C) for 0.5 h and then allow to recover at 22 °C for 180 min. Magenta arrowheads point toward GRP7 condensates. Condensate index are shown as means ± s.d..

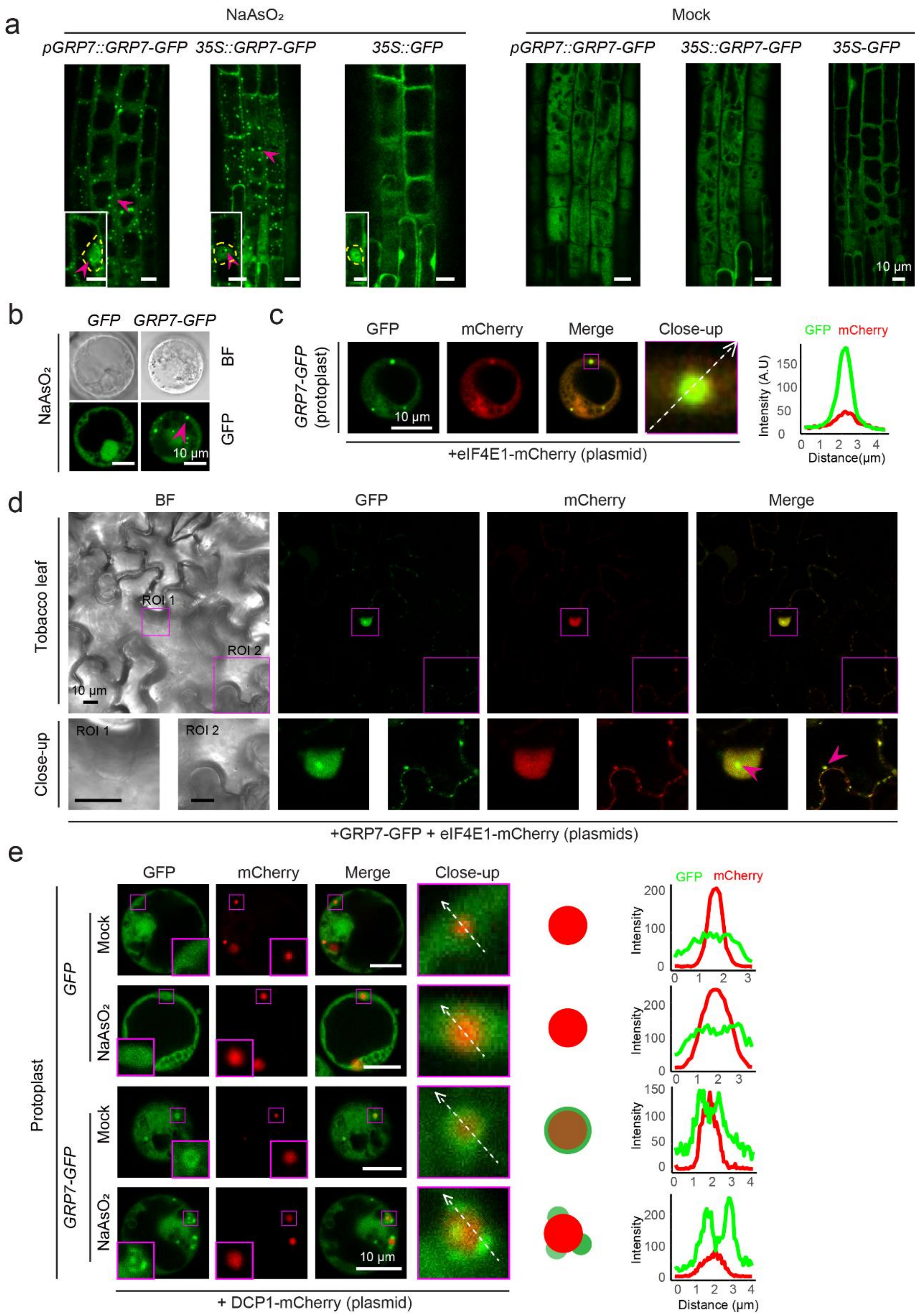


**Supplementary Fig. 3. Cytoplasmic GRP7 condensates are stress granules.**

**a**, GRP7 condensates in the cytoplasm and in the nucleus in the root cells of 5-d-old seedlings treated with 500 μM $NaAsO_2$ for 1h. Magenta arrowheads point to GRP7 condensates; yellow dashed lines enclose the nucleus. **b**, GRP7 condensates in root protoplasts induced by $NaAsO_2$. **c**, Co-localization of mCherry and GFP signals in protoplasts from *GRP7-GFP* roots expressing *eIF4E-mCherry*. The selected area (as indicated by the white dashed line in the close-up) was analysed for fluorescence intensity using FIJI, and the resulting plot profiles are shown at far right. **d**, Co-localization of mCherry and GFP signals in *N. benthamiana* leaves co-infiltrated with *GRP7-GFP* and *eIF4E-mCherry* constructs. **e**, Co-localization of mCherry and GFP signals in root protoplasts from *GRP7-GFP* seedlings expressing *DCP1-mCherry*. For **b**-**e**, protoplasts and *N. benthamiana* leaves were pretreated with 500 μM $NaAsO_2$ for 2 h before imaging. The diagrams represent the signal pattern of mCherry and GFP. The selected areas (as indicated by the white dashed lines in the close-up) were analysed for fluorescence intensity using FIJI, and the plot profiles are shown at far right. All fluorescence images were recorded using the same parameters by laser-scanning confocal microscopy. Two independent biological replicates were analysed, yielding similar results.

a

**Plate 1**

| | 1 | 2 | 3 | 4 | 5 | 6 | 7 | 8 | 9 | 10 | 11 | 12 |
|---|---|---|---|---|---|---|---|---|---|---|---|---|
| A | 0.1M Citric acid<br>pH 3.5,<br>20% PEG500 | 0.1M Citric acid<br>pH 3.5,<br>30% PEG500 | 0.1M Citric acid<br>pH 3.5,<br>10% PEG3350 | 0.1M Citric acid<br>pH 3.5,<br>20% PEG3350 | 0.1M Citric acid<br>pH 3.5,<br>10% PEG8000 | 0.1M Citric acid<br>pH 3.5,<br>20% PEG8000 | 0.1M Citric acid<br>pH 3.5,<br>10% ficoll400 | 0.1M Citric acid<br>pH 3.5,<br>20% ficoll400 | 0.1M Citric acid<br>pH 3.5,<br>10% Dextran10 | 0.1M Citric acid<br>pH 3.5,<br>20% Dextran10 | 0.1M Citric acid<br>pH 3.5,<br>10% Dextran40 | 0.1M Citric acid<br>pH 3.5,<br>20% Dextran40 |
| B | 0.1M Citric acid<br>pH 4.5,<br>20% PEG500 | 0.1M Citric acid<br>pH 4.5,<br>30% PEG500 | 0.1M Citric acid<br>pH 4.5,<br>10% PEG3350 | 0.1M Citric acid<br>pH 4.5,<br>20% PEG3350 | 0.1M Citric acid<br>pH 4.5,<br>10% PEG8000 | 0.1M Citric acid<br>pH 4.5,<br>20% PEG8000 | 0.1M Citric acid<br>pH 4.5,<br>10% ficoll400 | 0.1M Citric acid<br>pH 4.5,<br>20% ficoll400 | 0.1M Citric acid<br>pH 4.5,<br>10% Dextran10 | 0.1M Citric acid<br>pH 4.5,<br>20% Dextran10 | 0.1M Citric acid<br>pH 4.5,<br>10% Dextran40 | 0.1M Citric acid<br>pH 4.5,<br>20% Dextran40 |
| C | 0.1M Bis-Tris<br>pH 5.5,<br>20% PEG500 | 0.1M Bis-Tris<br>pH 5.5,<br>30% PEG500 | 0.1M Bis-Tris<br>pH 5.5,<br>10% PEG3350 | 0.1M Bis-Tris<br>pH 5.5,<br>20% PEG3350 | 0.1M Bis-Tris<br>pH 5.5,<br>10% PEG8000 | 0.1M Citric acid<br>pH 5.5,<br>20% PEG8000 | 0.1M Citric acid<br>pH 5.5,<br>10% ficoll400 | 0.1M Citric acid<br>pH 5.5,<br>20% ficoll400 | 0.1M Citric acid<br>pH 5.5,<br>10% Dextran10 | 0.1M Citric acid<br>pH 5.5,<br>20% Dextran10 | 0.1M Bis-Tris<br>pH 5.5,<br>10% Dextran40 | 0.1M Bis-Tris<br>pH 5.5,<br>20% Dextran40 |
| D | 0.1M Bis-Tris<br>pH 6.5,<br>20% PEG500 | 0.1M Bis-Tris<br>pH 6.5,<br>30% PEG500 | 0.1M Bis-Tris<br>pH 6.5,<br>10% PEG3350 | 0.1M Bis-Tris<br>pH 6.5,<br>20% PEG3350 | 0.1M Bis-Tris<br>pH 6.5,<br>10% PEG8000 | 0.1M Bis-Tris<br>pH 6.5,<br>20% PEG8000 | 0.1M Citric acid<br>pH 6.5,<br>10% ficoll400 | 0.1M Citric acid<br>pH 6.5,<br>20% ficoll400 | 0.1M Citric acid<br>pH 6.5,<br>10% Dextran10 | 0.1M Citric acid<br>pH 6.5,<br>20% Dextran10 | 0.1M Bis-Tris<br>pH 6.5,<br>10% Dextran40 | 0.1M Bis-Tris<br>pH 6.5,<br>20% Dextran40 |
| E | 0.1M HEPES<br>pH 7.5,<br>20% PEG500 | 0.1M HEPES<br>pH 7.5,<br>30% PEG500 | 0.1M HEPES<br>pH 7.5,<br>10% PEG3350 | 0.1M HEPES<br>pH 7.5,<br>20% PEG3350 | 0.1M HEPES<br>pH 7.5,<br>10% PEG8000 | 0.1M HEPES<br>pH 7.5,<br>20% PEG8000 | 0.1M Citric acid<br>pH 7.5,<br>10% ficoll400 | 0.1M Citric acid<br>pH 375,<br>20% ficoll400 | 0.1M Citric acid<br>pH 7.5,<br>10% Dextran10 | 0.1M Citric acid<br>pH 7.5,<br>20% Dextran10 | 0.1M HEPES<br>pH 7.5,<br>10% Dextran40 | 0.1M HEPES<br>pH 7.5,<br>20% Dextran40 |
| F | 0.1M Tris<br>pH 8.5,<br>20% PEG500 | 0.1M Tris<br>pH 8.5,<br>30% PEG500 | 0.1M Tris<br>pH 8.5,<br>10% PEG3350 | 0.1M Tris<br>pH 8.5,<br>20% PEG3350 | 0.1M Tris<br>pH 8.5,<br>10% PEG8000 | 0.1M Tris<br>pH 8.5,<br>20% PEG8000 | 0.1M Citric acid<br>pH 8.5,<br>10% ficoll400 | 0.1M Citric acid<br>pH 8.5,<br>20% ficoll400 | 0.1M Citric acid<br>pH 8.5,<br>10% Dextran10 | 0.1M Citric acid<br>pH 8.5,<br>20% Dextran10 | 0.1M Tris<br>pH 8.5,<br>10% Dextran40 | 0.1M Tris<br>pH 8.5,<br>20% Dextran40 |
| G | 0.1M Citric acid<br>pH 3.5,<br>10% Dextran70 | 0.1M Bis-Tris<br>pH 4.5,<br>10% Dextran70 | 0.1M HEPES<br>pH 5.5,<br>10% Dextran70 | 0.1M Citric acid<br>pH 6.5,<br>10% Dextran70 | 0.1M Bis-Tris<br>pH 7.5,<br>10% Dextran70 | 0.1M Tris<br>pH 8.5,<br>10% Dextran70 | 30% PEG500 | 20% PEG8000 | 20% Dextran40 | 0.1M Citric acid<br>pH 3.5 | 0.1M Bis-Tris<br>pH 5.5 | 0.1M HEPES<br>pH 7.5 |
| H | 0.1M Citric acid<br>pH 3.5,<br>20% Dextran70 | 0.1M Bis-Tris<br>pH 4.5,<br>20% Dextran70 | 0.1M HEPES<br>pH 5.5,<br>20% Dextran70 | 0.1M Citric acid<br>pH 6.5,<br>20% Dextran70 | 0.1M Bis-Tris<br>pH 7.5,<br>20% Dextran70 | 0.1M Tris<br>pH 8.5,<br>20% Dextran70 | 20% PEG3350 | 20% Dextran10 | 20% Dextran70 | 0.1M Citric acid<br>pH 4.5 | 0.1M Citric acid<br>pH 6.5 | 0.1M Citric acid<br>pH 8.5 |

b

**Plate 2**

| | 1 | 2 | 3 | 4 | 5 | 6 | 7 | 8 | 9 | 10 | 11 | 12 |
|---|---|---|---|---|---|---|---|---|---|---|---|---|
| A | 0.1M Citric acid<br>pH 3.5,<br>1M NaCl | 0.1M Citric acid<br>pH 3.5,<br>3M NaCl | 0.1M Citric acid<br>pH 3.5,<br>1M ammonium<br>sulfate | 0.1M Citric acid<br>pH 3.5,<br>3M ammonium<br>sulfate | 100μM CR7 | 400μM CR7 | 0.75M<br>Lithium chloride | 0.2M<br>Zinc acetate | 0.2M<br>Zinc acetate<br>20% PEG3350 | 0.2M<br>Calcium<br>chloride | 0.2M<br>Calcium<br>chloride<br>20% PEG3350 | |
| B | 0.1M Citric acid<br>pH 4.5<br>1M NaCl | 0.1M Citric acid<br>pH 4.5,<br>3M NaCl | 0.1M Citric acid<br>pH 4.5,<br>1M ammonium<br>sulfate | 0.1M Citric acid<br>pH 4.5,<br>3M ammonium<br>sulfate | 100μM CR20 | 400μM CR20 | 0.75M<br>Lithium chloride<br>20% PE3350 | 0.5M<br>Zinc acetate | 0.5M<br>Zinc acetate<br>20% PEG3350 | 0.5M<br>Calcium<br>chloride | 0.5M<br>Calcium<br>chloride<br>20% PEG3350 | |
| C | 0.1M Citric acid<br>pH 5.5<br>1M NaCl | 0.1M Citric acid<br>pH 5.5,<br>3M NaCl | 0.1M Citric acid<br>pH 5.5,<br>1M ammonium<br>sulfate | 0.1M Citric acid<br>pH 5.5,<br>3M ammonium<br>sulfate | 100μM heparin | 100μM heparin<br>20% PEG3350 | 100μM heparin<br>20% PEG8000 | 0.75M<br>Zinc acetate | 0.75M<br>Zinc acetate<br>20% PEG3350 | 0.75M<br>Calcium<br>chloride | 0.75M<br>Calcium<br>chloride<br>20% PEG3350 | |
| D | 0.1M Bis-Tris<br>pH 6.5<br>1M NaCl | 0.1M Citric acid<br>pH 6.5,<br>3M NaCl | 0.1M Citric acid<br>pH 6.5,<br>1M ammonium<br>sulfate | 0.1M Citric acid<br>pH 6.5,<br>3M ammonium<br>sulfate | 400μM heparin | 400μM heparin<br>20% PEG3350 | 400μM heparin<br>20% PEG8000 | 0.2M<br>Zinc acetate | 0.2M<br>Magnesium<br>chloride<br>20% PEG3350 | 0.2M<br>Magnesium<br>sulfate | 0.2M<br>Magnesium<br>sulfate<br>20% PEG3350 | |
| E | 0.1M HEPES<br>pH 7.5<br>1M NaCl | 0.1M Citric acid<br>pH 7.5,<br>3M NaCl | 0.1M Citric acid<br>pH 7.5,<br>1M ammonium<br>sulfate | 0.1M Citric acid<br>pH 7.5,<br>3M ammonium<br>sulfate | 1000μM<br>heparin | 1000μM<br>heparin<br>20% PEG3350 | 1000μM<br>heparin<br>20% PEG8000 | 0.2M<br>Zinc acetate | 0.5M<br>Magnesium<br>chloride<br>20% PEG3350 | 0.5M<br>Magnesium<br>sulfate | 0.5M<br>Magnesium<br>sulfate<br>20% PEG3350 | |
| F | 0.1M HEPES<br>pH 8.5<br>1M NaCl | 0.1M Citric acid<br>pH 8.5,<br>3M NaCl | 0.1M Citric acid<br>pH 8.5,<br>1M ammonium<br>sulfate | 0.1M Citric acid<br>pH 8.5,<br>3M ammonium<br>sulfate | 2μg/μL polyU | 2μg/μL polyU<br>20% PEG3350 | 2μg/μL polyU<br>20% PEG8000 | 0.2M<br>Zinc acetate | 0.75M<br>Magnesium<br>chloride<br>20% PEG3350 | 0.75M<br>Magnesium<br>sulfate | 0.75M<br>Magnesium<br>sulfate<br>20% PEG3350 | |
| G | 0.2M<br>Potassium<br>acetate | 0.5M<br>Potassium<br>acetate | 0.75M<br>Potassium<br>acetate | 0.2M<br>Sodium acetate | 0.5M<br>Sodium acetate | 0.75M<br>Sodium acetate | 0.2M<br>ammonium<br>chloride | 0.5M<br>ammonium<br>chloride | 0.75M<br>ammonium<br>chloride | 0.2M<br>Lithium<br>chloride | 0.5M<br>Lithium<br>chloride | |
| H | 0.2M<br>Potassium<br>acetate<br>20% PEG3350 | 0.5M<br>Potassium<br>acetate<br>20% PEG3350 | 0.75M<br>Potassium<br>acetate<br>20% PEG3350 | 0.2M<br>Sodium acetate<br>20% PEG3350 | 0.5M<br>Sodium acetate<br>20% PEG3350 | 0.75M<br>Sodium acetate<br>20% PEG3350 | 0.2M<br>ammonium<br>chloride<br>20% PEG3350 | 0.5M<br>ammonium<br>chloride<br>20% PEG3350 | 0.75M<br>ammonium<br>chloride<br>20% PEG3350 | 0.2M<br>Lithium<br>chloride<br>20% PEG3350 | 0.5M<br>Lithium<br>chloride<br>20% PEG3350 | |

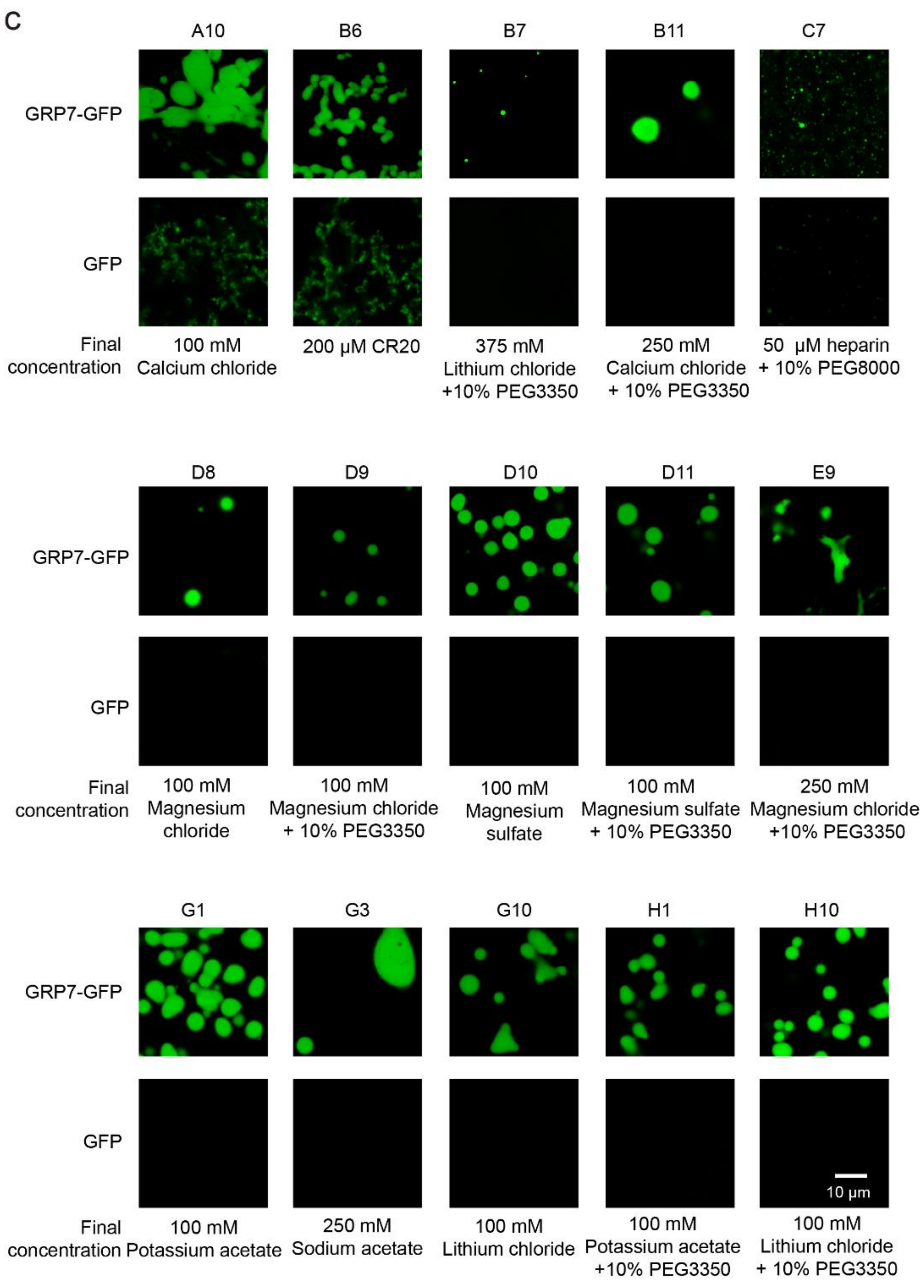


**Supplementary Fig. 4. Screening of GRP7 phase separation conditions.**

**a** and **b**, Compositions of the solutions in the wells of the HiPSS kit. The solutions induced GRP7-GFP phase separation are highlighted in green. **c**, Representative fluorescence images of phase-separated GRP7-GFP for specific wells of the HiPSS kit (Plate 2). Each well contains 50 μM GRP7-GFP or GFP.

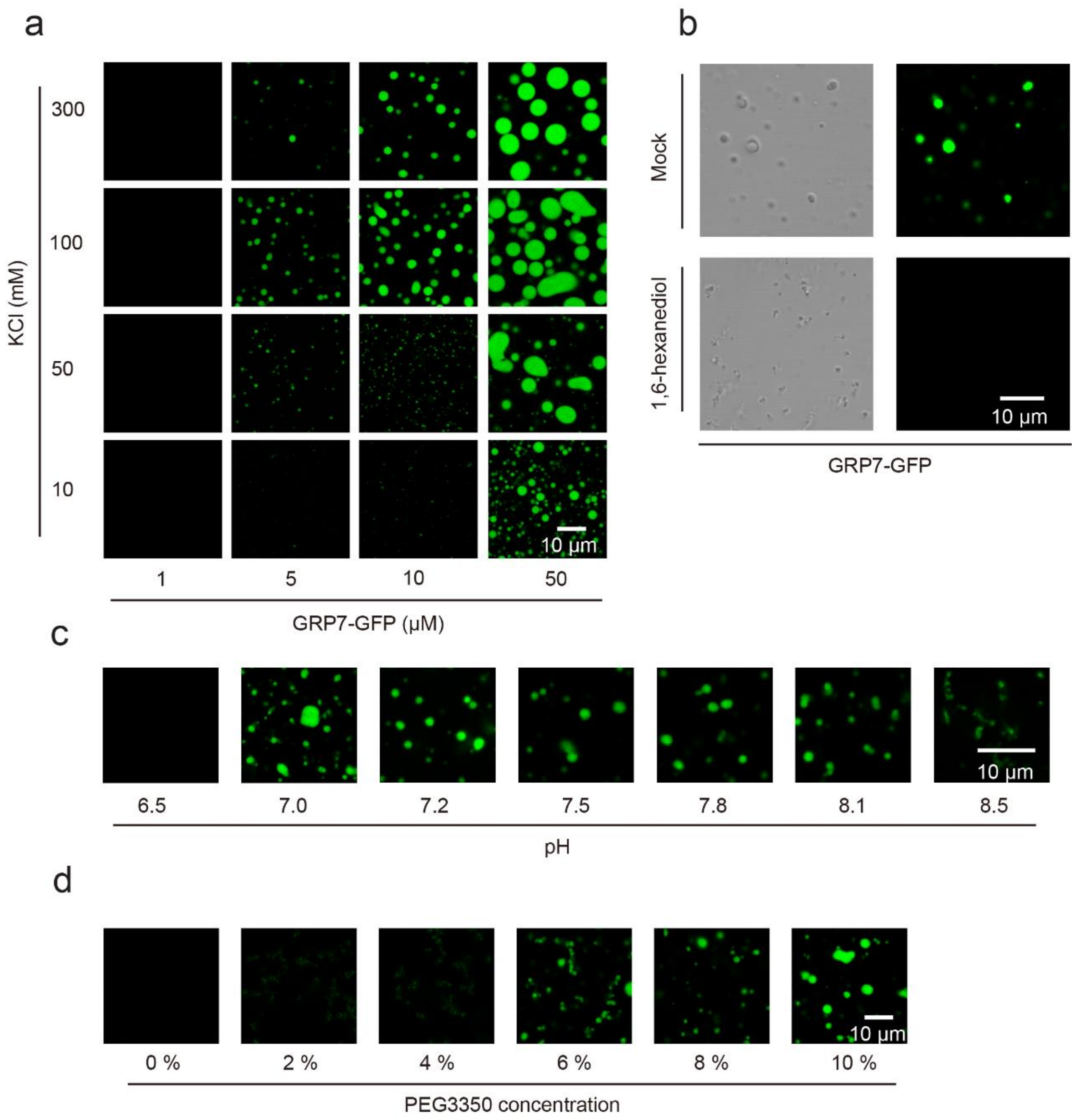


**Supplementary Fig. 5. GRP7 phase separation characteristics.**

**a**, Representative images related to the GRP7-GFP phase-separation diagram. **b**, GRP7-GFP (10 μM) droplets are disrupted by 5% (v/v) 1,6-hexanediol. **c**, GRP7-GFP (10 μM) droplet formation at different pH values. **d**, Effect of PEG concentration on GRP7-GFP droplet formation. In **c**,**d**, GRP7-GFP (10 μM) was mixed with 100 mM KCl and 10% (w/v) PEG3350.

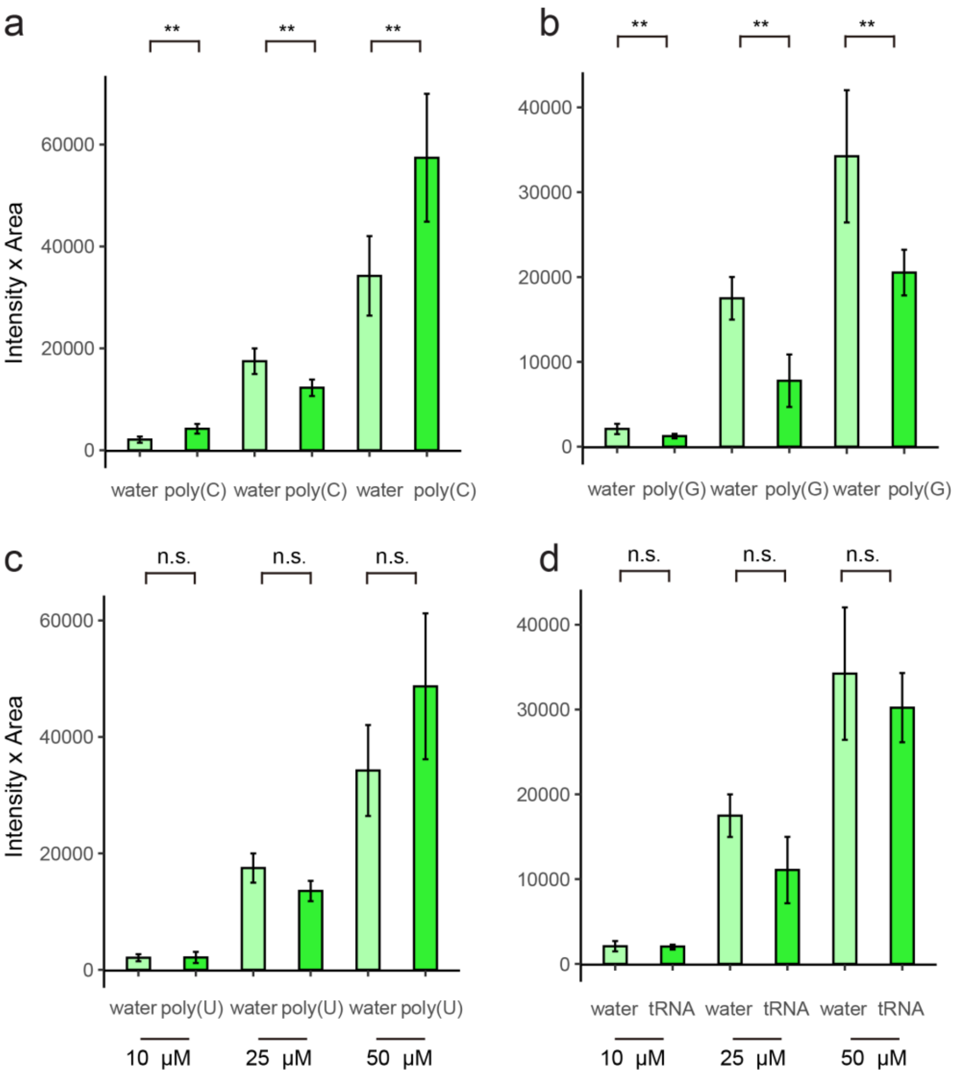


**Supplementary Fig. 6. Effect of poly(C), poly(G), poly(U) and tRNA on GRP7 phase separation *in vitro*.**

**a**-**d**. Sum of the mean fluorescence intensity × area for all droplets per field of view from three images. Data are shown as means ± s.d.; **P* < 0.05, ***P* < 0.01, n.s., not significant by one-way ANOVA. **a**, poly(C); **b**, poly(G); **c**, poly(U); **d**, tRNA.

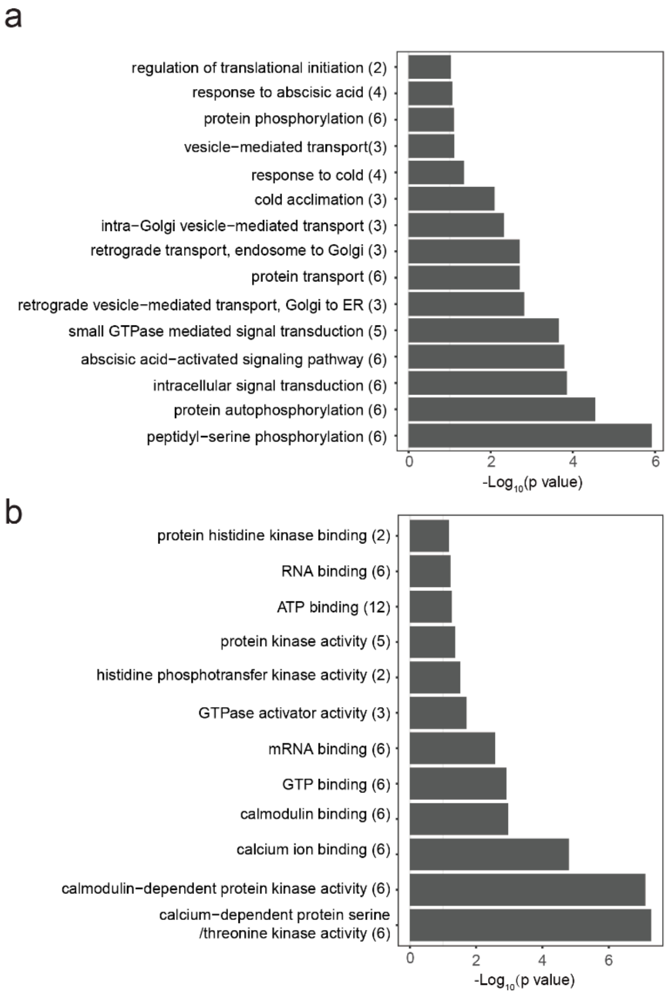


**Supplementary Fig. 7. Gene Ontology term analysis of proteins identified in cytoplasmic GRP7 condensates.**

**a**-**b**, The list of 47 proteins identified in in cytoplasmic GRP7-GFP condensates was submitted to the ShinyGO v0.60 Gene Ontology Enrichment Analysis tool (http://bioinformatics.sdstate.edu/go/) using default parameters. Fisher's exact test was applied to identify most enriched GO terms in Biological Process (**a**) and Molecular Function (**b**).

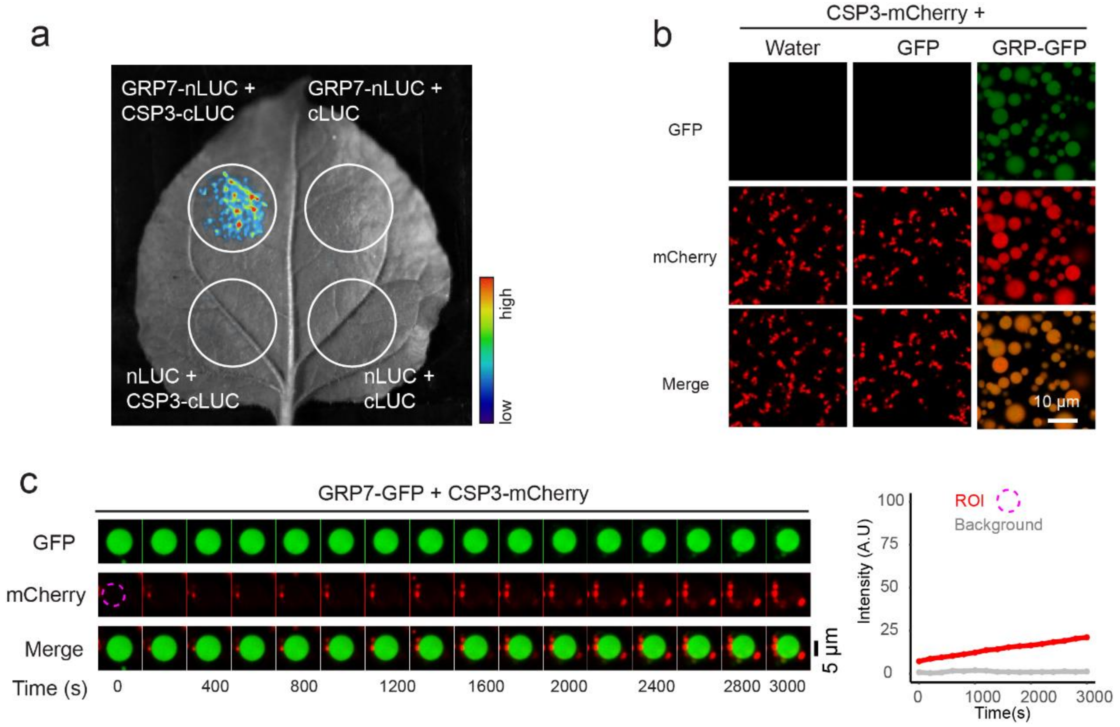


#### Supplementary Fig. 8. GRP7 phase separation recruits CSP3.

**a**, Split luciferase complementation assays showing the interaction between GRP7 and CSP3. The indicated constructs were co-infiltrated in *N. benthamiana* leaves through Agrobacterium-mediated infiltration. Luciferase activity was determined 48 h after infiltration. **b**, GRP7-GFP droplets recruited CSP3-mCherry. Liquid droplets formed after 50 μM GRP7-GFP was mixed with 50 μM CSP3-mCherry and incubated for 10 min at room temperature. **c**, Phase-separated GRP7-GFP droplets recruit the free form of CSP3-mCherry. A phase-separated GRP7-GFP solution was mixed with a solution of CSP3-mCherry, and images were acquired immediately. Regions of interest are highlighted by the dashed magenta circle. Right, changes in the intensity of CSP3-mCherry over time.

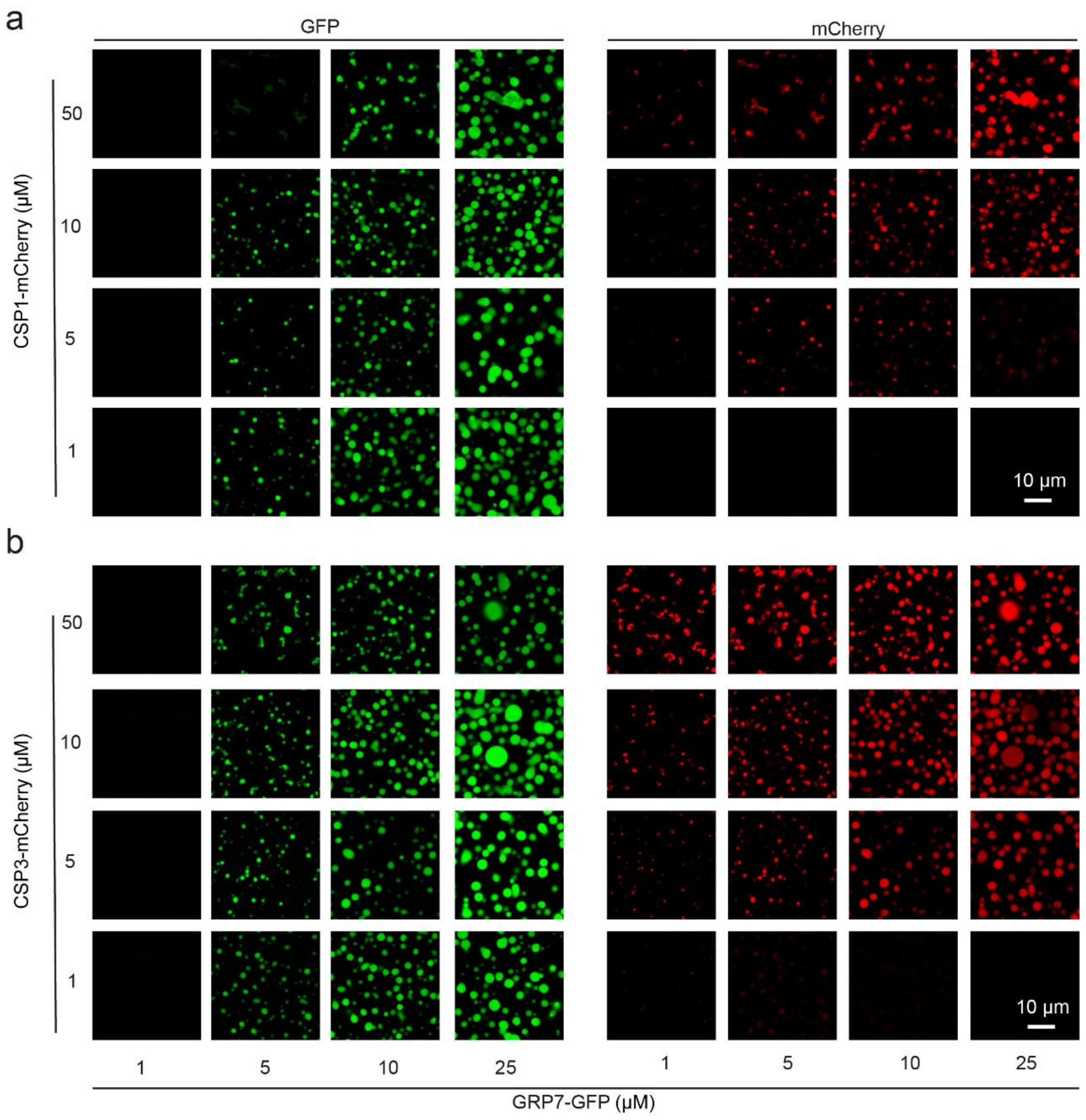


**Supplementary Fig. 9. Phase-separated GRP7-GRP incorporates CSP1- and CSP3-mCherry.**

**a**,**b**, Representative images related to the GRP7-GFP phase separation diagram for various concentrations of CSP1-mCherry (**a**) and CSP3-mCherry (**b**).

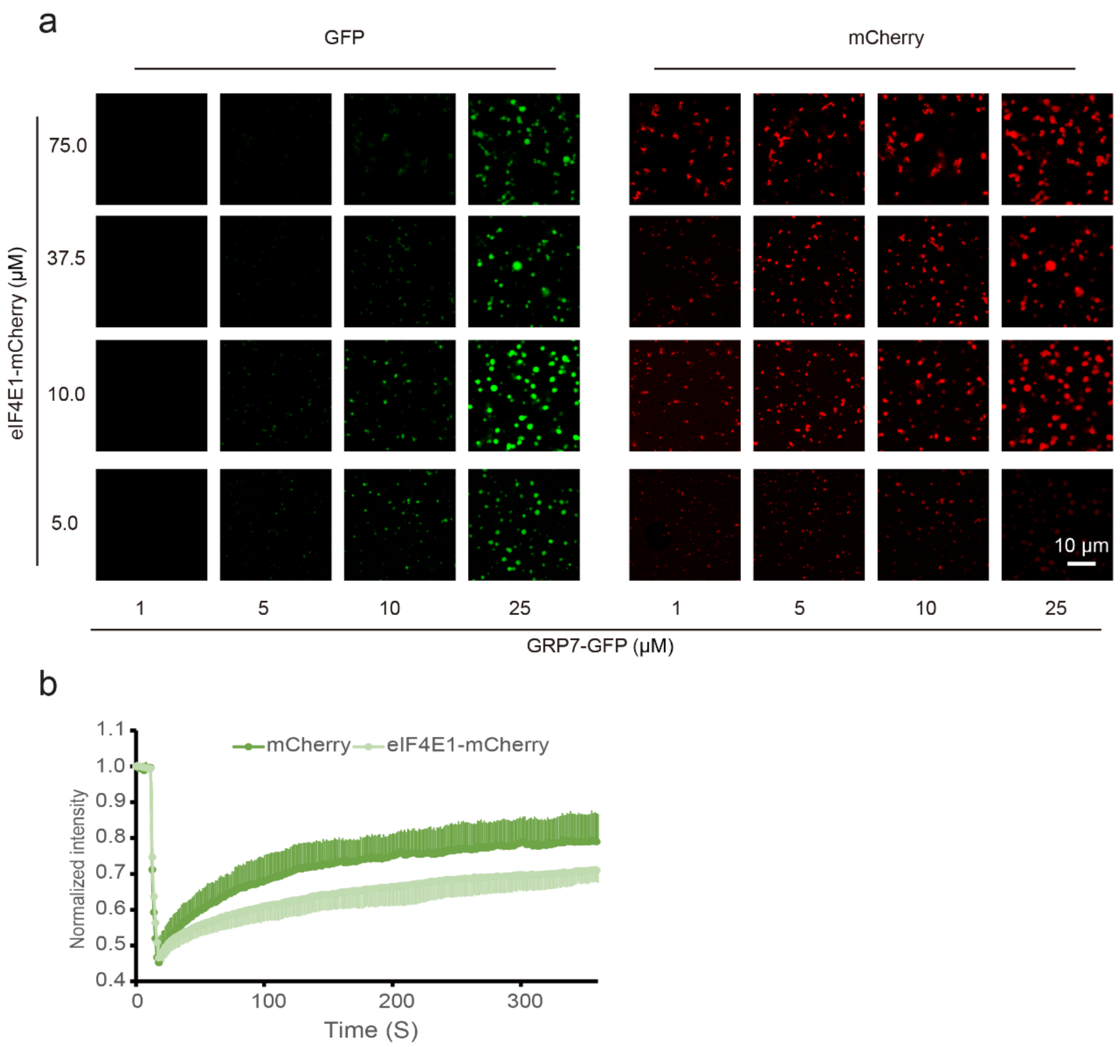


**Supplementary Fig. 10. eIF4E1-mCherry influences GRP7-GFP phase separation.**

a, Representative images related to the GRP7-GFP phase separation diagram for various concentrations of eIF4E1-mCherry. b, Effect of eIF4E1-mCherry on FRAP of GRP7-GFP.

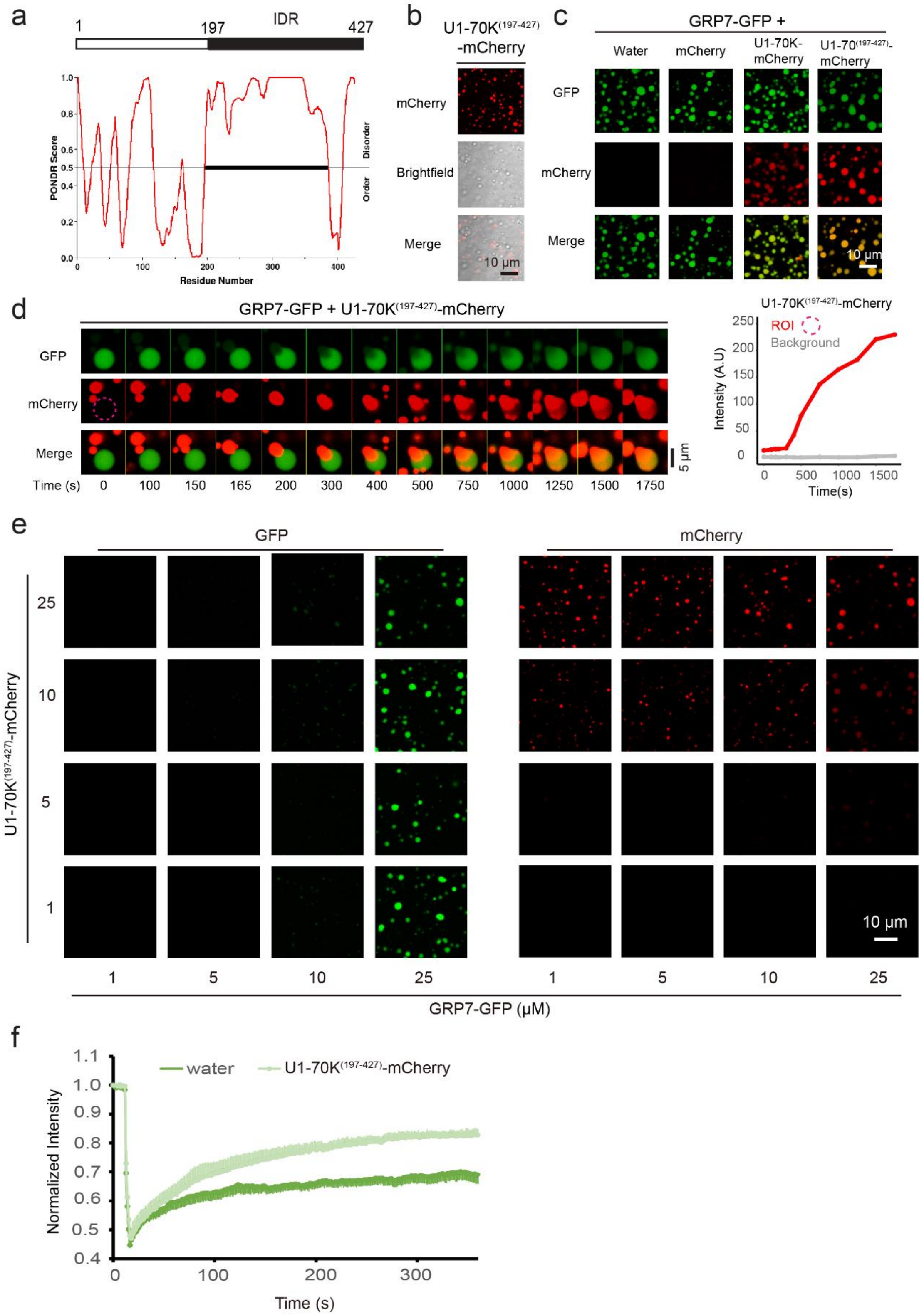


**Supplementary Fig. 11. Phase-separated GRP7-GRP incorporates truncated U1-70K-mCherry.**

**a**, Intrinsic disorder tendency of U1-70K, as predicted by the PONDR tool. The most disordered regions in these proteins are marked with a black line. **b**, U1-70K$^{(197-427)}$-mCherry (50 μM) phase separated in 10% PEG 3350 and 100 mM KCl. **c**, GRP7-GFP droplets recruited U1-70K-mCherry and U1-70K$^{(197-}$

$^{427)}$-mCherry. Liquid droplets formed after mixing 50 µM GRP7-GFP with 15 µM eIF4E1-mCherry, U1-70K-mCherry and U1-70K$^{(197-427)}$-mCherry and incubating for 10 min at room temperature. **d**, Phase-separated GRP7-GFP droplets recruit the phase-separated form of U1-70K$^{(197-427)}$-mCherry. A phase-separated GRP7-GFP solution was mixed with a solution of phase-separated U1-70K$^{(197-427)}$-mCherry, and images were acquired immediately. Region of interest is highlighted by the dashed magenta circle. The right panel indicates changes in the intensity of U1-70K$^{(197-427)}$-mCherry over time. **e**, Representative images related to the GRP7-GFP phase-separation diagram for various concentrations of U1-70K$^{(197-427)}$-mCherry. **f**, Effect of U1-70K$^{(197-427)}$-mCherry on FRAP of GRP7-GFP.

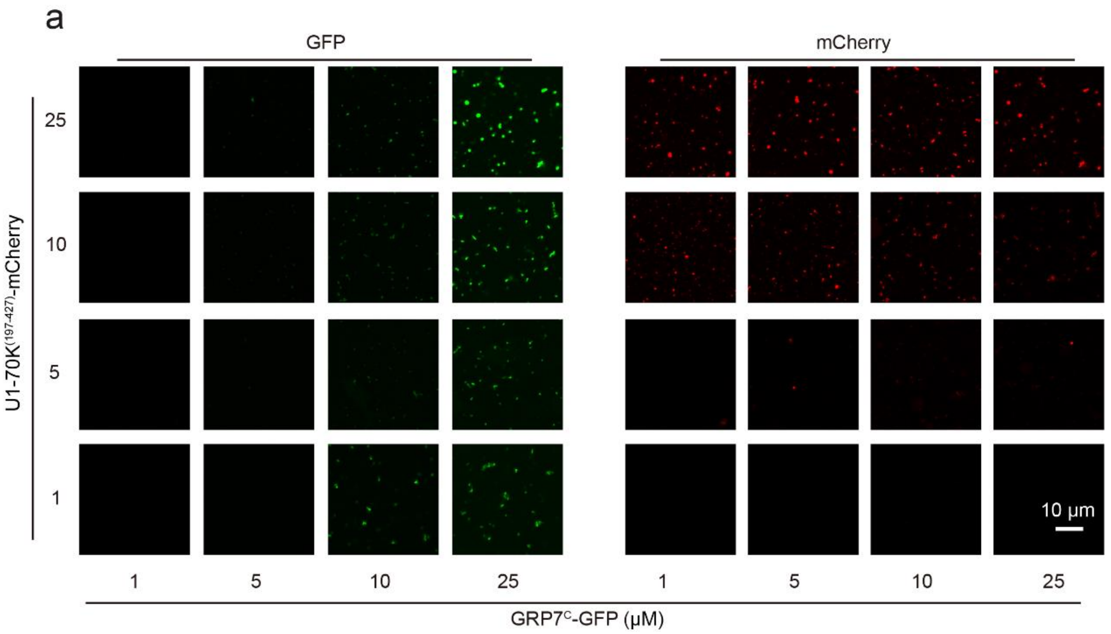


**Supplementary Fig. 12. Phase-separated GRP7C-GRP incorporates truncated U1-70K-mCherry.**

**a**, Representative images related to the GRP7C-GFP phase separation diagram for various concentrations of U1-70K$^{(197-427)}$-mCherry.

**Supplementary Movie 1. GRP7 condensates in Arabidopsis root cells. Related to Fig. 2.**

**Supplementary Movie 2. Two GRP7 condensates fusing in a root cell. Related to Figure 2.**

**Supplementary Movie 3. Trajectory of GRP7 condensates in a root cell. Related to Fig. 2.**

**Supplementary Movie 4. 3D view of two fusing GRP7 condensates during an *in vitro* phase-separation assay. Related to Fig. 3.**

**Supplementary Movie 5. Phase-separated U1-70K$^{(197\text{-}427)}$-mCherry is incorporated in to phase-separated GRP7-GFP droplets. Related to Supplementary Fig. 11.**

**Supplementary Movie 6. Free eIF4E-mCherry is incorporated into phase-separated GRP7-GFP droplets. Related to Fig. 6.**

**Supplementary Table 1. Proteins identified in CO-IP MS.**

**Supplementary Table 2. Primers used in this study.**
